# ESCAPE: assigning site-specific activity to covalent ligands in cells by prime editing

**DOI:** 10.64898/2026.07.28.741261

**Authors:** Jason E. Tse, William R. Brothers, Rachel E. Hayward, Sabrina Barbas, Kai Sheng, Kathryn R. Spencer, Kristen E. DeMeester, Evert Njomen, James R. Williamson, David R. Liu, Bruno Melillo, Gene W. Yeo, Benjamin F. Cravatt, Haoxin Li

## Abstract

Chemical proteomics has identified covalent ligands targeting cysteine residues across many hundreds of human proteins. The functional effects of these liganding events, however, remain challenging to assign at scale. Here we describe ESCAPE (Endogenous Site-specific Competition Assays using Prime Editors), a platform for the site-resolved functional analysis of covalent ligands in cells. In this method, cysteine-to-serine substitutions are generated by prime editing to abrogate covalent ligand-protein interactions, and the impact of these edits on ligand-induced cellular phenotypes is quantified through allele frequency-based resistance scores. Applied to ligandable cysteines mapped by activity-based protein profiling in 50+ proteins, ESCAPE identified multiple covalent ligand-protein interactions that impair cancer cell growth, including azetidine butynamides that target a non-orthosteric cysteine in the RNA helicase DDX49 to disrupt 18S rRNA processing, 40S ribosome assembly, and protein synthesis. ESCAPE thus provides a scalable framework for the functional characterization of covalent ligands targeting structurally and mechanistically diverse proteins.

## Introduction

Small molecules serve as critical tools for studying the biochemical and cellular functions of proteins and represent a major category of therapeutics^1^. Most human proteins, however, lack selective chemical probes. To address this gap, several innovative technologies for small-molecule screening have been introduced that include fragment-based ligand discovery^2^, DNA-encoded libraries^3,4^, and activity-based protein profiling (ABPP)^5–7^.

As a chemical proteomic approach, ABPP has the advantage of mapping small molecule-protein interactions directly in living cells^8,9^. When applied to electrophilic compounds using a mass spectrometry (MS) readout, ABPP can further identify the precise residues on proteins that react with small molecules^10–15^. This approach has led to the discovery of first-in-class covalent ligands for a diverse range of proteins, including historically challenging classes such as transcription factors, adaptors, E3 ligases, and RNA-binding proteins^16–23^.

Many of the covalent liganding events mapped by ABPP occur with cysteine residues located at non-orthosteric sites on proteins and determining the functional impact of these small molecule-protein interactions can be an arduous process for several reasons. First, in contrast to small molecules targeting orthosteric sites (e.g., enzyme active sites), where the functional impact (inhibition) can be inferred, non-orthosteric ligands can produce a wide range of modulatory effects on proteins, including antagonism, (neo)agonism, and silent outcomes^24,25^. There are even instances where the same non-orthosteric pocket can support both agonistic and antagonistic functional effects based on subtle changes in ligand structure^26–29^. Second, the proteins engaged by covalent ligands originate from a wide range of mechanistically distinct classes, each of which may require its own specialized biochemical or cellular assay for understanding ligand effects on protein activity. Finally, the covalent hit ligands discovered by ABPP have not yet been optimized for potency or proteomic selectivity and therefore have the potential to produce cellular effects reflecting complex, multi-target activities. While these challenges have been addressed for individual covalent ligand-protein pairs by leveraging, for instance, structurally related inactive control compounds (e.g., enantiomers of hit ligands) and ligand-resistant cysteine mutant recombinant proteins^11,18,19,30^, a systematic way to functionally characterize many covalent ligand-protein interactions in parallel is lacking.

Here, we describe a scalable method termed ESCAPE (Endogenous Site-specific Competition Assays using Prime Editors) for site-resolved functional profiling of covalent ligands in human cells. ESCAPE first leverages prime editing^31^-mediated cysteine-to-serine (C-to-S) substitutions to prevent covalent ligand engagement at defined sites in the endogenous proteome. Then, following covalent compound treatment, ESCAPE determines the contribution of individual ligandable sites to cellular phenotypes measured through quantifying wild-type versus C-to-S allele frequencies. Applying ESCAPE to 50+ covalent ligand-protein interactions mapped by ABPP, we identify several small molecules that perturb cancer cell growth through covalent modification of individual cysteine residues in the proteome. These anti-proliferative compounds include an azetidine butynamide MY-45B that stereoselectively reacts with a non-orthosteric cysteine (C258) in the RNA helicase DDX49, leading to impairments in 18S rRNA processing, ribosome biogenesis, and protein synthesis in cancer cells. Interestingly, ESCAPE also identified analogs of MY-45B that act as silent DDX49 ligands in cells, underscoring the potential for this method to distinguish functional from silent small molecules binding at the same protein site.

## Results

### Workflow and benchmarking of ESCAPE

ESCAPE requires a quantifiable phenotype for comparing the functional effects of covalent compounds in cells endogenously expressing WT vs compound-resistant (C-to-S) mutant forms of proteins. A review of ABPP data for stereochemically defined electrophilic small molecules (stereoprobes)^11,16,30,32^ identified stereoprobe-liganded cysteines in many proteins that support the growth of cancer cells as determined by genome-wide CRISPR screens described in the Cancer Dependency Map (DepMap)^33,34^. We accordingly reasoned that, for these ligandable cancer dependency proteins, cell proliferation could serve as a phenotypic measure of covalent compound effects, where an active compound would be expected to impair the growth of cells expressing the WT, but not compound-resistant C-to-S mutant form of its target protein(s) (**Fig. 1a**). In brief, prime editors, representing a Cas9 nickase fused to a reverse transcriptase, were used to install precise genomic edits encoding C-to-S mutations specified by pegRNAs^35–39^. Edited cell populations (minipools of WT protein-expressing and prime-edited C-to-S protein-expressing cells) were then generated in 96-well format and treated with DMSO or covalent stereoprobes for 5 days, followed by targeted genomic PCR and sequencing (**Fig. 1a**). A resistance score was defined as the ratio of the acquired serine allele frequency in compound-treated versus DMSO control conditions, providing a quantitative measure of the functional contribution of site-specific covalent engagement of cysteine residues in cancer dependency proteins to cell proliferation (**Fig. 1a**).

**Fig. 1.**
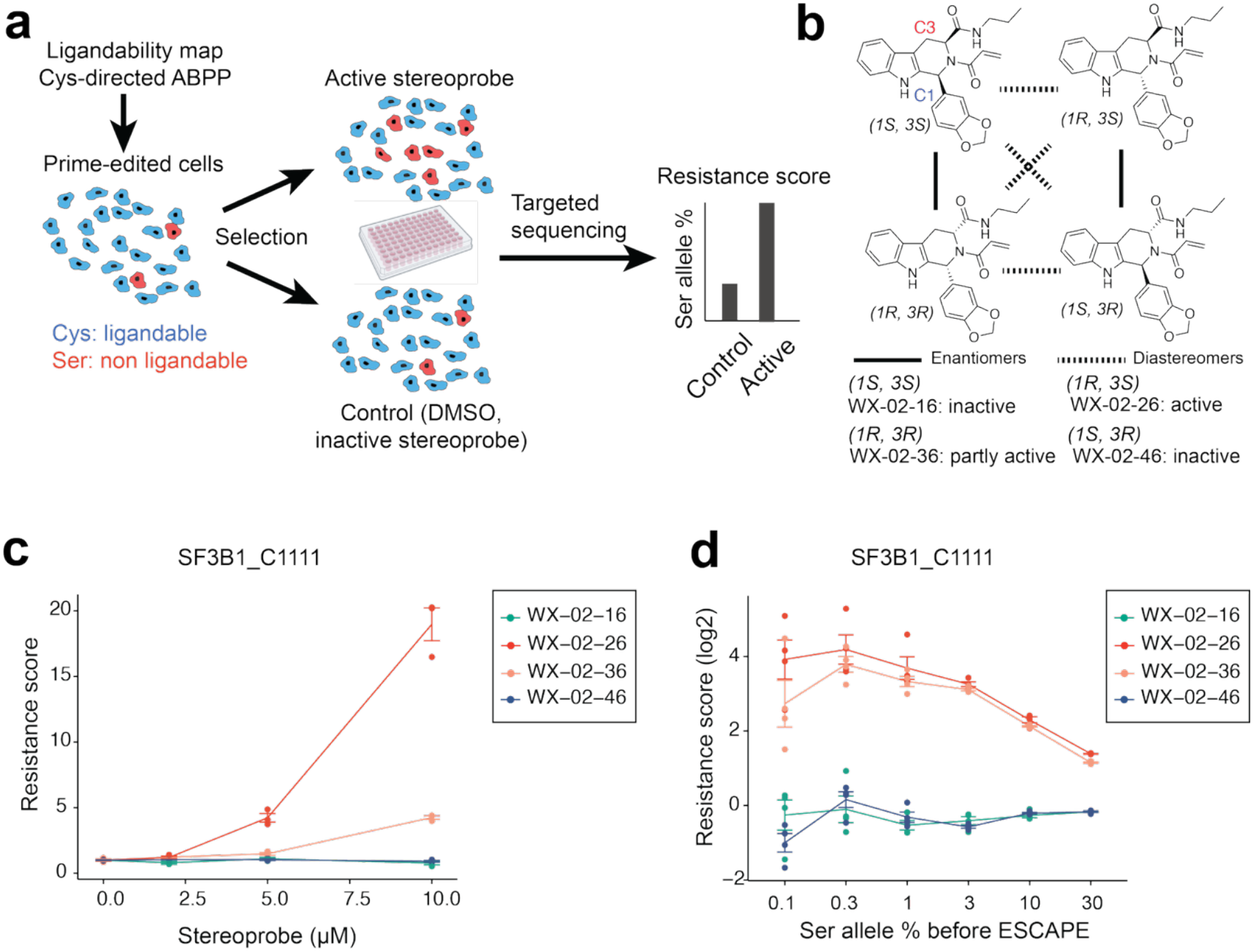
Workflow and benchmarking of ESCAPE. **a**, Schematic illustrating the ESCAPE workflow. Ligandable cysteines (Cys) identified by ABPP in cancer dependency proteins are converted to covalent ligand-resistant serines (Ser) by prime editing. Cells are then treated with covalent ligands or DMSO for 5 days, followed by targeted genomic sequencing. Resistance scores are defined as the ratio of Ser allele frequency in probe-treated versus DMSO control conditions. **b**, Structures of tryptoline acrylamide stereoprobes displaying the indicated engagement profiles for cysteine-1111 (C1111) of the common essential protein SF3B1 used for benchmarking ESCAPE. See ref.^11^ and **Extended Data Fig. 1a** for tryptoline acrylamide engagement profiles for SF3B1. **c**, ESCAPE assay performed in HCT116 cells harboring the C1111S-SF3B1 edit treated with the tryptoline acrylamide stereoprobes from **b** at the indicated concentrations. Data are average values ± SEM from 3 independent experiments. **d**, Assessing the performance of ESCAPE at different editing frequencies achieved by mixing different fractions of C1111S-SF3B1 edited and parental (wild type-SF3B1) HCT116 cells (X axis). Cells were then treated with 10 µM of the indicated stereoprobes for 5 days, and resistance scores were calculated (Y axis). The C1111S-SF3B1 cells used in this study represent a clonal line generated as described in the **Methods** section. Data are average values ± SEM from 4 independent experiments.

To benchmark ESCAPE, we evaluated tryptoline acrylamide stereoprobes targeting C1111 of the essential splicing factor SF3B1 (**Fig. 1b**). Previous ABPP experiments have identified (1*R*, 3*S*) tryptoline acrylamides (e.g., WX-02-26) that enantioselectively engage SF3B1, as well as a similar, but weaker enantioselective liganding of SF3B1 by (1*R*, 3*R*) tryptoline acrylamides (e.g., WX-02-36)^11,16^ (**Fig. 1b** and **Extended Data Fig. 1a**). Functional studies further verified that SF3B1-directed tryptoline acrylamides produce enantioselective and site-specific impairments in RNA splicing and cancer cell growth^16^. Mirroring the ligandability profile of SF3B1, ESCAPE experiments revealed that WX-02-26, and to a lesser extent WX-02-36, produced concentration-dependent resistance scores in a pool of wild type and prime-edited C1111S-SF3B1-edited HCT116 cells, while the inactive enantiomeric compounds (WX-02-46 and WX-02-16, respectively) showed negligible effects (**Fig. 1c**). ESCAPE experiments generating the C1111A-SF3B1 allele yielded comparable resistance scores in cells treated with the active ligands (**Extended Data Fig. 1b**). These data thus provided initial evidence to support that ESCAPE can quantify the functional impact of covalent ligands targeting an essential protein in cancer cells.

We next evaluated the sensitivity of ESCAPE by mixing C1111S-SF3B1-edited HCT116 cells with wild-type cells at defined ratios. Notably, ESCAPE quantified strong resistance scores even when the initial edited S allele frequency was as low as 0.1% (**Fig. 1d**). Conversely, and as expected, the magnitude of resistance score began to decrease as the edited S allele frequency rose above 10% (**Fig. 1d**). These results highlight that ESCAPE can be applied to cells bearing very low frequency gene edits and also help to define a dynamic range of editing that is most suitable for measuring resistance to the anti-proliferative effects of covalent ligands.

Together, our initial studies with covalent stereoprobes targeting SF3B1 supported the potential for ESCAPE to serve as a quantitative and sensitive platform for characterizing the anti-proliferative effects of covalent small molecule-protein interactions in cells.

### ESCAPE identifies covalent liganding events that impair cancer cell growth

We applied ESCAPE to a panel of 60 stereoprobe-liganded cysteines mostly found at non-orthosteric sites on proteins from diverse structural and functional classes (**Fig. 2a** and **Table 1**). The proteins harboring liganded cysteines exhibited a range of essentiality for cancer cell growth based on the DepMap^33,34^, including 38 common essential and 12 non-essential proteins that were profiled in the colon cancer cell line HCT116, as well as 10 proteins showing selective dependency profiles that were examined in context-matched dependent cell lines (**Fig. 2b** and **Supplementary Table 1**). Importantly, the stereoprobes evaluated by ESCAPE mostly represented hit compounds not yet optimized for high proteomic selectivity and thus provided a valuable test for method performance at an early stage in the assessment of the bioactivity of covalent ligands targeting sites on proteins of unclear functionality. With this consideration in mind, we tested stereoprobes at concentrations expected to produce at least 60% cysteine engagement (5-20 µM (**Table 1)**, as inferred from previous ABPP experiments^11,16,30,32,40^, and **Supplementary Table 2**), while also limiting the potential for gross toxicity caused by general electrophile stress.

**Fig. 2.**
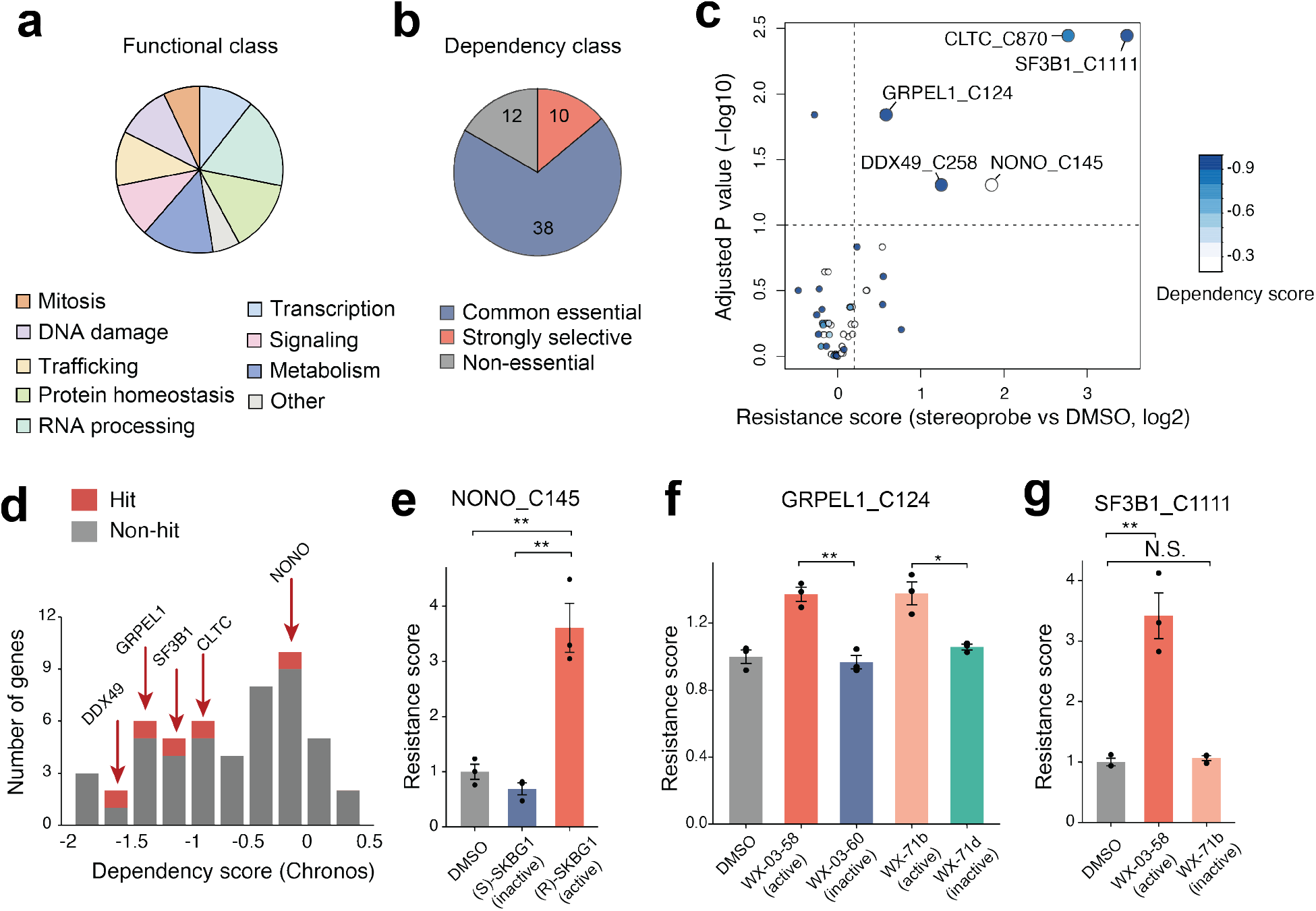
ESCAPE identifies covalent liganding events that impair cancer cell growth. **a**, Pie chart showing the distribution of protein functional classes evaluated by ESCAPE. **b**, Pie chart showing the proportion of proteins evaluated by ESCAPE from the indicated DepMap dependency classes (common essential, strongly selective dependency, or non-essential). See **Table 1** for list of stereoprobe-protein interactions tested by ESCAPE. **c**, Volcano plot showing the resistance scores and adjusted P values for the covalent ligand-protein interactions assayed by ESCAPE in HCT116 cells treated with active stereoprobes vs DMSO control. Prime-edited HCT116 cells harboring Cys-to-Ser point mutations for each protein target were treated with respective stereoprobe ligands (5-15 µM, see **Table 1**) or DMSO for 5 days, followed by targeted genomic PCR (see **Supplementary Table 1** and **Methods** section). Dots for covalent ligand-protein interactions defined as hits in the ESCAPE assay (resistance scores (log2) > 0.2 with adjusted P values (-log10) > 1) are enlarged for visualization purposes. **d**, Histogram showing the distribution of Chronos gene dependency scores in HCT116 cells from the DepMap for the corresponding proteins with ligandable cysteines assayed by ESCAPE. Hits identified in (**c**) are highlighted in red. **e**, ESCAPE results for NONO_C145 evaluated with the active covalent ligand (*R*)-SKBG1 and the inactive enantiomer (*S*)-SKBG1^40^ (5 μM, 5 days) in HCT116 cells. **f**, ESCAPE results for GRPEL1_C124 evaluated with the indicated active tryptoline acrylamide stereoprobes (5 μM, 5 days) and their respective inactive enantiomers^11,32^ in HCT116 cells. **g**, ESCAPE results for SF3B1_C1111 evaluated with the indicated tryptoline acrylamide stereoprobes^11,32^ (5 μM, 5 days) in HCT116 cells. For all ESCAPE results, data are average values ± SEM from 3 independent experiments. All P values were calculated using two-sided Student’s t test. N.S., not significant, * p<0.05, **p<0.01, ***p<0.001.

**Table 1.** Covalent stereoprobe-protein interactions evaluated by ESCAPE. For each interaction, the table lists the liganded protein and cysteine, the active and inactive stereoprobe pair used in ESCAPE assay, the stereoprobe concentration used, and the cell line selected based on the target protein’s genetic dependency profile in the DepMap^33,34^. Types of gene dependencies: CE, Common Essential; SS, Strongly Selective; NE, Non-Essential.

| Liganded Protein_Cysteine | DepMap | Active Stereoprobe | Inactive Stereoprobe | References | Ligand (μM) | Cancer Cell line |
| --- | --- | --- | --- | --- | --- | --- |
| ATRIP_C577 | CE | WX-03-348 | WX-03-346 | Ref.<sup>11</sup> | 10 | HCT116 |
| BLK_C319 | NE | WX-03-340 | WX-03-338 | Ref.<sup>11</sup> | 10 | HCT116 |
| CACTIN_C699 | CE | WX-03-347 | WX-03-349 | Ref.<sup>11</sup> | 10 | HCT116 |
| CLTC_C870 | CE | WX-02-34 | WX-02-44 | This paper, ref.<sup>30</sup> | 10 | HCT116 |
| CNOT1_C1932 | CE | WX-71d | WX-71b | Ref.<sup>32</sup> | 10 | HCT116 |
| COG1_C87 | CE | WX-03-349 | WX-03-347 | Ref.<sup>11</sup> | 10 | HCT116 |
| DCTN4_C258 | CE | WX-02-16 | WX-02-46 | Ref.<sup>11</sup> | 10 | HCT116 |
| DDA1_C25 | CE | WX-01-06 | WX-01-08 | Ref.<sup>11</sup> | 10 | HCT116 |
| DDX49_C258 | CE | MY-45B | MY-45A | This paper, ref.<sup>16</sup> | 10 | HCT116 |
| ELL_C14 | CE | WX-02-46 | WX-02-26 | Ref.<sup>11</sup> | 10 | HCT116 |
| ELMO1_C438 | NE | WX-02-16 | WX-02-36 | Ref.<sup>11</sup> | 10 | HCT116 |
| ELP2_C746 | CE | WX-03-63 | WX-03-61 | This paper | 10 | HCT116 |
| EP400_C1820 | CE | WX-02-46 | WX-02-26 | Ref.<sup>11</sup> | 10 | HCT116 |
| ESYT2_C181 | NE | WX-02-26 | WX-02-46 | Ref.<sup>11</sup> | 10 | HCT116 |
| EXOSC9_C9 | CE | WX-01-12 | WX-01-10 | Ref.<sup>11</sup> | 10 | HCT116 |
| FBL_C268 | CE | WX-72d | WX-72b | Ref.<sup>32</sup> | 10 | HCT116 |
| GPHN_C419 | NE | WX-03-63 | WX-03-61 | This paper | 10 | HCT116 |
| GRPEL1_C124 | CE | WX-71b | WX-71d | Ref.<sup>32</sup> | 10 | HCT116 |
| GSDMD_C268 | NE | WX-01-12 | WX-01-10 | Ref.<sup>11</sup> | 10 | HCT116 |
| GTF3C3_C607 | CE | WX-72d | WX-72b | Ref.<sup>32</sup> | 10 | HCT116 |
| INTS13_C349 | CE | WX-03-64 | WX-03-62 | This paper | 10 | HCT116 |
| INTS8_C491 | CE | MY-45B | MY-45A | This paper, ref.<sup>16</sup> | 10 | HCT116 |
| LTN1_C723 | NE | WX-03-60 | WX-03-58 | Ref.<sup>11</sup> | 10 | HCT116 |
| MAST3_C541 | NE | WX-03-339 | WX-03-341 | Ref.<sup>11</sup> | 10 | HCT116 |
| MCCC2_C216 | NE | WX-03-338 | WX-03-340 | Ref.<sup>11</sup> | 10 | HCT116 |
| MMS19_C819 | CE | WX-03-349 | WX-03-347 | Ref.<sup>11</sup> | 10 | HCT116 |
| MMS22L_C466 | CE | WX-02-26 | WX-02-46 | Ref.<sup>11</sup> | 10 | HCT116 |
| MRPL39_C133 | CE | WX-01-12 | WX-01-10 | Ref.<sup>11</sup> | 10 | HCT116 |
| MVD_C386 | CE | WX-01-11 | WX-01-09 | Ref.<sup>11</sup> | 10 | HCT116 |
| NCAPD2_C766 | CE | WX-02-46 | WX-02-26 | Ref.<sup>11</sup> | 10 | HCT116 |
| NCAPD2_C767 | CE | WX-02-46 | WX-02-26 | Ref.<sup>11</sup> | 10 | HCT116 |
| NCAPD3_C631 | CE | WX-02-36 | WX-02-16 | Ref.<sup>11</sup> | 10 | HCT116 |
| NONO_C145 | SS | (R)-SKBG-1 | (S)-SKBG-1 | Ref.<sup>40</sup> | 5 | HCT116 |
| NOP16_C36 | CE | WX-02-14 | WX-02-34 | This paper, ref.<sup>30</sup> | 10 | HCT116 |
| NPEPPS_C339 | CE | WX-01-05 | WX-01-07 | Ref.<sup>11</sup> | 10 | HCT116 |
| NT5DC1_C119 | NE | WX-03-348 | WX-03-346 | Ref.<sup>11</sup> | 10 | HCT116 |
| NUDT16L1_C88 | NE | WX-02-16 | WX-02-36 | Ref.<sup>11</sup> | 10 | HCT116 |
| NUP98_C1312 | CE | WX-03-60 | WX-03-58 | Ref.<sup>11</sup> | 10 | HCT116 |
| PDPK1_C21 | CE | WX-03-60 | WX-03-58 | Ref.<sup>11</sup> | 10 | HCT116 |
| PDPK1_C23 | CE | WX-03-60 | WX-03-58 | Ref.<sup>11</sup> | 10 | HCT116 |
| POLE_C2148 | CE | WX-72b | WX-72d | Ref.<sup>32</sup> | 10 | HCT116 |
| SARS2_C64 | CE | WX-03-60 | WX-03-58 | Ref.<sup>11</sup> | 10 | HCT116 |
| SARS2_C66 | CE | WX-03-60 | WX-03-58 | Ref.<sup>11</sup> | 10 | HCT116 |
| SF3B1_C1111S | CE | WX-02-26 | WX-02-46 | Ref.<sup>11</sup> | 10 | HCT116 |
| SMG5_C109 | CE | WX-03-340 | WX-03-338 | Ref.<sup>11</sup> | 10 | HCT116 |
| SNW1_C250 | CE | WX-01-06 | WX-01-08 | Ref.<sup>11</sup> | 10 | HCT116 |
| SRBD1_C294 | CE | WX-01-12 | WX-01-10 | Ref.<sup>11</sup> | 10 | HCT116 |
| TBC1D2B_C726 | NE | WX-02-36 | WX-02-16 | Ref.<sup>11</sup> | 10 | HCT116 |
| TEC_C449 | NE | WX-03-58 | WX-03-60 | Ref.<sup>11</sup> | 10 | HCT116 |
| TRRAP_C620 | CE | WX-01-06 | WX-01-08 | Ref.<sup>11</sup> | 10 | HCT116 |
| VPS18_C806 | CE | WX-03-57 | WX-03-59 | Ref.<sup>11</sup> | 10 | HCT116 |
| CDS2_C286 | SS | MY-45A | MY-45B | This paper, ref.<sup>16</sup> | 10 | OVCAR8 |
| CHUK_C406 | SS | WX-02-308 | WX-02-326 | This paper, ref.<sup>84</sup> | 15 | KMS11 |
| CYB5B_C119 | SS | WX-03-349 | WX-03-347 | Ref.<sup>11</sup> | 10 | K562 |
| EEFSEC_C442 | SS | WX-02-36 | WX-02-16 | Ref.<sup>11</sup> | 20 | TC71 |
| LIG1_C895 | SS | WX-64d | WX-64b | Ref.<sup>32</sup> | 10 | OVCAR8 |
| PDS5A_C532 | SS | WX-03-60 | WX-03-58 | Ref.<sup>11</sup> | 10 | NUGC3 |
| RPP25_C70 | SS | WX-03-340 | WX-03-338 | Ref.<sup>11</sup> | 15 | CALU6 |
| SOS1_C405 | SS | WX-02-37 | WX-02-17 | Ref.<sup>30</sup> | 10 | K562 |
| STAMBP_C264 | SS | WX-03-341 | WX-03-339 | Ref.<sup>11</sup> | 15 | 22RV1 |

Using a threshold of >15% increase in resistance score (active stereoprobe_vs_DMSO control) and a false discovery rate (FDR) <10%, we identified five functional stereoprobe-cysteine interactions, including the positive control WX-02-26:SF3B1_C1111 liganding event (**Fig. 2c**). Four of the five hits corresponded to essential proteins as defined by the DepMap with corresponding gene dependency scores ranging from -0.89 to -1.72 (**Fig. 2d**), whereas the single non-essential protein hit represented the RNA-binding protein NONO. Four of the five hits corresponded to common essential proteins as defined by the DepMap (**Fig. 2d**), whereas the fifth hit, the RNA-binding protein NONO, showed a strongly selective dependency profile in DepMap but was not essential in HCT116 cells, the cell line used in our screen (**Table 1**). In previous work, we have shown that the NONO_C145 ligand (*R*)-SKBG1, but not the inactive enantiomer (*S*)-SKBG1, stabilizes NONO binding to mRNAs in cells leading to a gain-of-toxicity (rather than loss-of-function) outcome that likely explains the disconnect between the DepMap gene dependency (**Fig. 2d**) and ESCAPE resistance score values (**Fig. 2e**) for this protein^40^. We observed similar ESCAPE results when comparing active stereoprobes to their inactive enantiomers, with one additional functional stereoprobe-protein interaction being identified (WX-02-14:NOP16_C36; **Extended Data Fig. 2a**). This interaction just missed the cutoff for being defined as a hit in the active stereoprobe-vs-DMSO comparison (resistance score =1.46, adjusted P value = 0.246).

One of the essential protein hits from ESCAPE – GRPEL1 – functions as a nucleotide exchange factor for the mitochondrial Hsp70 chaperone HSPA9, and we have recently found that stereoprobe ligands targeting C124 of GRPEL1 disrupt interactions with HSPA9, leading to defects in mitochondrial protein import and respiration^32^. Interestingly, one of the tryptoline acrylamide stereoprobes WX-03-58 that produced a stereoselective ESCAPE signal with GRPEL1_C124 (**Fig. 2f**) also engages SF3B1_C1111 and likewise generated a strong ESCAPE signal with this protein (**Fig. 2g**). Additionally, a more optimized tryptoline acrylamide ligand for GRPEL1 – WX-71b – that lacks SF3B1 cross-reactivity ^32^ produced a similar ESCAPE score for GRPEL1_C124 as WX-03-58 (**Fig. 2f**), while showing no effect with SF3B1_C1111 (**Fig. 2g**). We interpret these results to indicate that ESCAPE is capable of deconvoluting the functional contributions of individual liganding events distributed across multiple essential proteins targeted by the same compound.

Taken together, the results of our screen revealed that ESCAPE can identify established functional covalent ligand-protein interactions that perturb cancer cell growth, including ligands that act by a gain-of-function (rather than loss-of-function) mechanism and that simultaneously engage more than one essential protein in cells. We next set out to investigate stereoprobes targeting DDX49_C258 (**Fig. 2c**), which represented a newly discovered functional liganding event mapped by ESCAPE.

### Stereoprobes targeting DDX49 perturb cancer cell growth

The ESCAPE screen yielded a positive resistance score for azetidine butynamide MY-45B in cells harboring a DDX49_C258S edit (**Fig. 2c** and **Fig. 3a, b**). Previous cysteine-directed ABPP experiments had shown the enantioselective engagement of DDX49_C258 by MY-45B^16^ (**Fig. 3c** and **Extended Data Fig. 3a**), and we confirmed and extended these findings by protein-directed ABPP (**Extended Data Fig. 3b** and **Supplementary Table 2**). These ABPP experiments also identified a limited number of additional cysteines and proteins that cross-reacted with MY-45B (**Extended Data Fig. 3a, b**). Gel-ABPP experiments performed with alkyne analogs of MY-45B and its enantiomer MY-45A (WX-04-500 and WX-04-499, respectively) verified enantioselective engagement of recombinant WT-DDX49, but not a C258S-DDX49 mutant protein, stably expressed in HCT116 cells (**Fig. 3d** and **Extended Data Fig. 3c**). We also found that MY-45B enantioselectively reacted with purified WT-DDX49, but not the C258S-DDX49 mutant protein by intact MS analysis (**Extended Data Fig. 3d, e**).

**Fig. 3.**
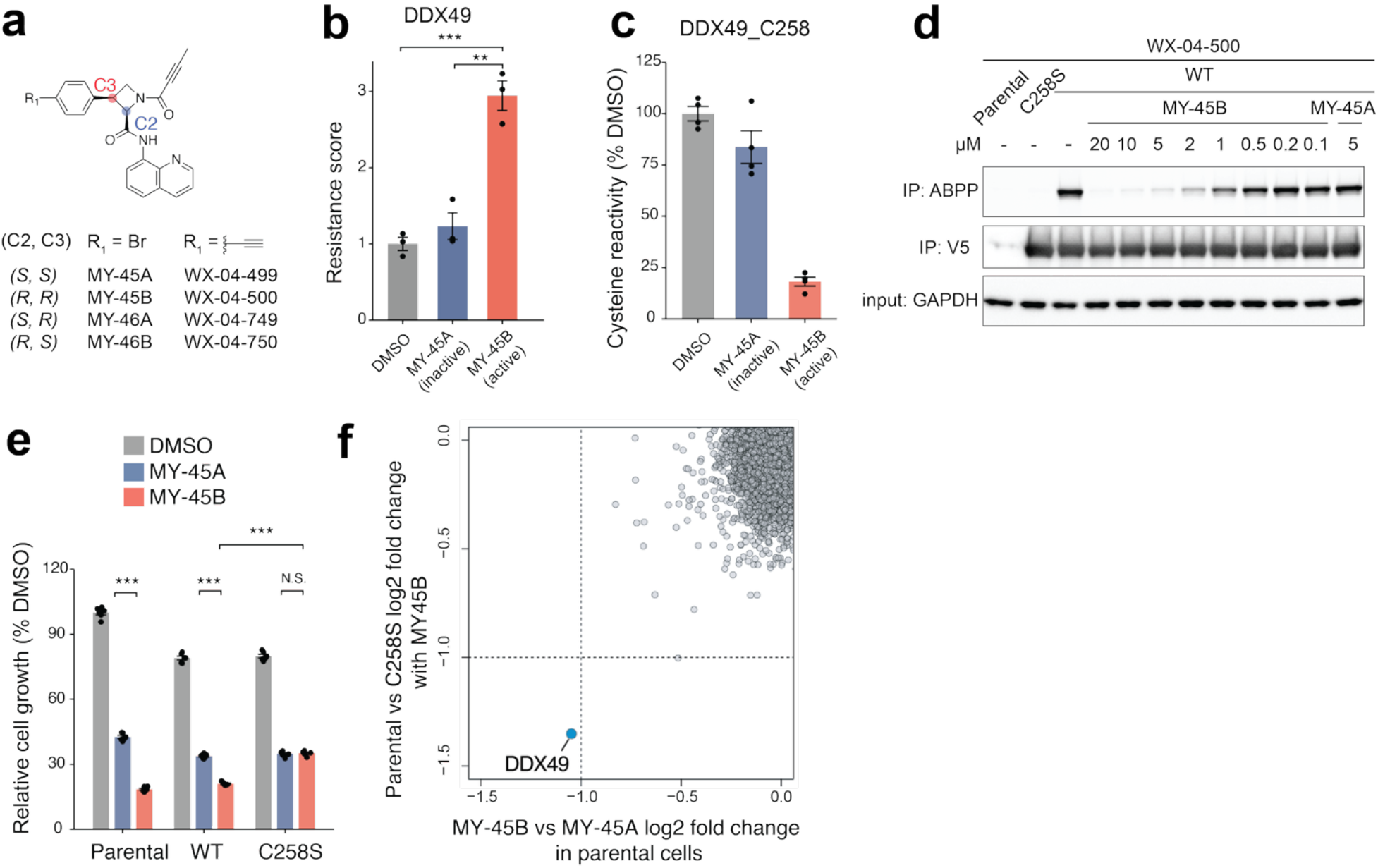
Stereoprobes targeting DDX49 perturb cancer cell growth. **a**, Structures of azetidine butynamide stereoprobes, including the DDX49_C258 ligand MY-45B^16^, and the respective alkyne analogs with indicated absolute stereochemistry. The stereochemical configuration shown in the figure corresponds to that of MY-45B and its alkyne-containing analog WX-04-500. **b**, ESCAPE results for DDX49_C258 evaluated with the active stereoprobe MY-45B and the inactive enantiomer MY-45A (5 μM, 5 days) in HCT116. Data are average values ± SEM from 3 independent experiments. **c**, Cysteine-directed ABPP data derived from ref.^16^ showing stereoselective liganding of DDX49_C258 by MY-45B, but not MY-45A in 22Rv1 cells (5 μM, 3 h). Data represent average values ± SEM of 4 independent experiments. **d**, Gel-ABPP data showing engagement of recombinant WT-DDX49, but not a C258S-DDX49 mutant, by WX-04-500 and the concentration-dependent blockade of the WX-04-500-WT-DDX49 interaction by MY-45B, but not MY-45A. Parental, WT-DDX49-, and C258S-DDX49-expressing HCT116 cells were treated with MY-45A or MY-45B (3 h), followed by alkyne WX-04-500 (2 μM, 1 h) and analysis by gel-ABPP. Data are from a single experiment representative of 2 independent experiments. **e**, Relative growth effects of the indicated stereoprobes (5 μM) in parental, WT-DDX49-, and C258S-DDX49-expressing HCT116 cells. Cells were treated with compounds for 4 days prior to CellTiter-Glo measurement. Data are average values ± SEM from 6 independent experiments and normalized to DMSO-treated parental cells. **f**, Scatter plot showing effects of gene knockdown on the relative cell growth from genome-wide CRISPRi screen performed in HCT116 cells treated with MY-45A or MY-45B (5 μM, 12 days of probe treatment). X axis: comparison between MY-45B vs MY-45A. Y axis: comparison between MY-45B-treated parental cells vs MY-45B-treated C258S-DDX49-expressing cells. All P values were calculated using two-sided Student’s t test. N.S., not significant, * p<0.05, **p<0.01, ***p<0.001.

We confirmed that MY-45B exhibited greater anti-proliferative activity than MY-45A in both parental and WT-DDX49-expressing HCT116 cells, but not in C258S-DDX49-expressing HCT116 cells (**Fig. 3e** and **Extended Data Fig. 3f**). We also performed a genome-wide CRISPRi screen (57,050 sgRNAs targeting 18,901 genes^41^) in HCT116 cells, which identified DDX49 knockdown as the strongest sensitizing genetic modifier for the stereoselective (compared to MY-45A) or site-specific (compared to C258S-DDX49-expressing cells) effects of MY-45B (**Fig. 3f** and **Supplementary Table 3**). We confirmed the sensitizing effects of DDX49 knockdown on the enantioselective anti-proliferative activity of MY-45B with three independent sgRNAs (**Extended Data Fig. 3g, h**). These results are consistent with a loss-of-function mechanism for MY-45B due to inhibition of DDX49 activity, with the CRISPRi-dependent decrease in DDX49 abundance further compromising cancer cell proliferation in the presence of the compound.

### MY-45B impairs ribosome biogenesis and protein synthesis in cancer cells

DDX49 is a member of the DEAD-box family of ATP-dependent RNA helicases (**Fig. 4a**) and resides in the nucleolus of cells, where it plays a role in ribosomal RNA (rRNA) maturation and ribosome biogenesis^42^. Structural modeling of DDX49 using AlphaFold2^43^, combined with a comparison to the structure of the homologous helicase DDX19B bound to RNA^44^, placed the liganded C258 residue at a non-orthosteric site distal to the ATP-binding pocket and proximal to the predicted RNA-binding cleft (**Fig. 4b**). To better understand how MY-45B reactivity with C258 affects the function of DDX49, we examined the rRNA and ribosome content of cancer cells.

**Fig. 4.**
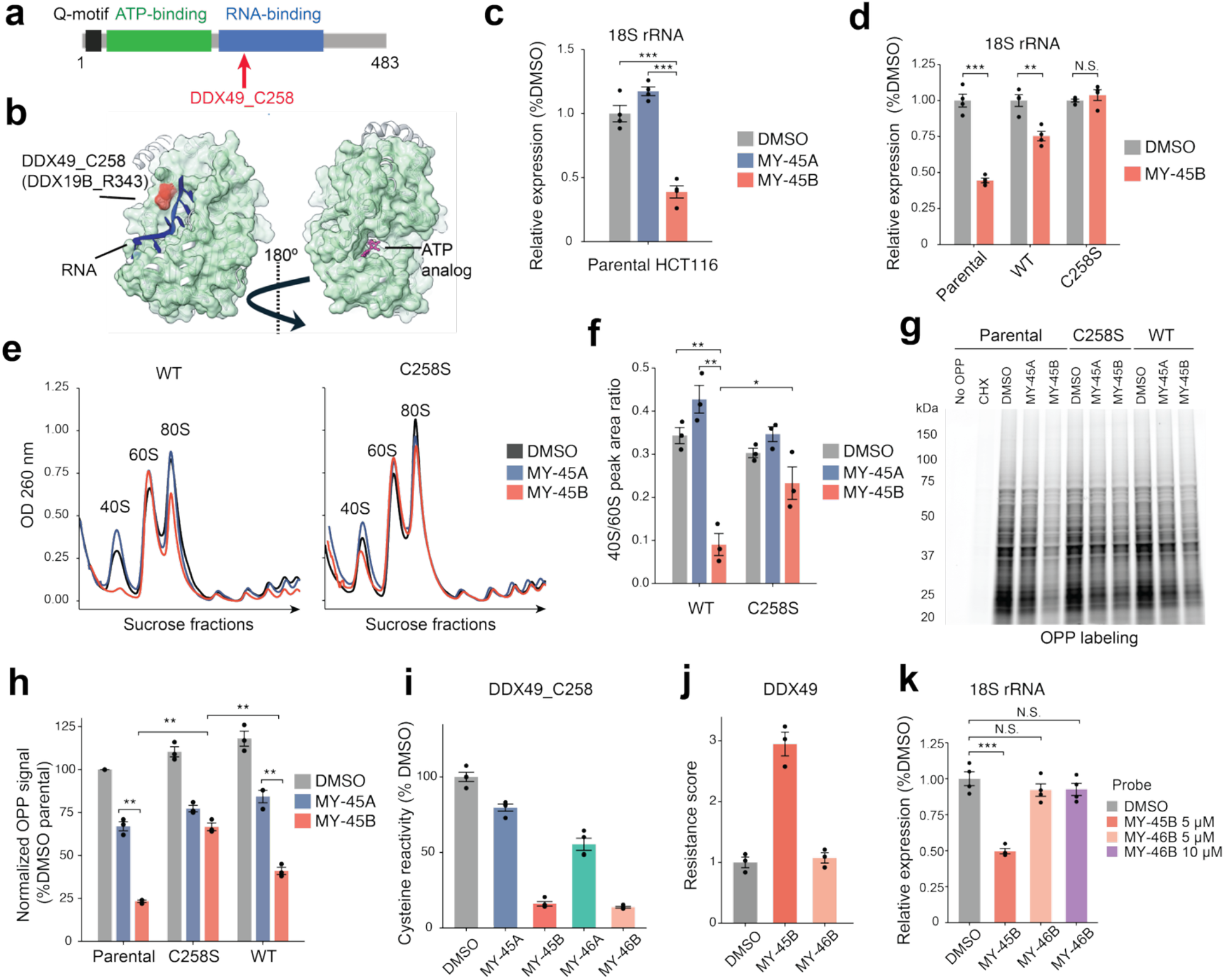
MY-45B impairs ribosome biogenesis and protein synthesis in cancer cells. **a**, Cartoon showing the domain architecture of DDX49 with C258 highlighted in red. **b**, Surface representation of the structure of the DDX49 homologous protein DDX19B bound to RNA (PDB: 3G0H) overlaid with an AF2-generated DDX49 structure (AF-Q9Y6V7, white ribbon). The MY-45B-liganded C258 of DDX49 and the analogous R343 residue in DDX19B are highlighted in red. **c**, 18S rRNA signals in parental HCT116 cells treated with MY-45A or MY-45B (5 μM, 48 h) as measured by qPCR. Data are average values ± SEM from 4 independent experiments. **d**, 18S rRNA signals in parental, WT-DDX49-expressing, and C258S-DDX49-expressing HCT116 cells treated with MY-45B (5 μM, 48 h) or DMSO as measured by qPCR. Data are average values ± SEM from 4 independent experiments and normalized internally to its own DMSO control. **e**, Polysome profiling of WT-DDX49-expressing or C258S-DDX49-expressing HCT116 cells treated with MY-45A or MY-45B (5 μM, 48 h) or DMSO. Results are from a single experiment representative of 3 independent experiments. **f**, Quantification of polysome profiling data for 40S and 60S peak areas for experiment shown in **e** and replicates. Data are average values ± SEM from 3 independent experiments. **g**, SDS-PAGE analysis of nascent protein synthesis in parental, WT-DDX49-expressing, and C258S-DDX49-expressing HCT116 cells treated with MY-45A or MY-45B (5 μM, 48 h) or DMSO followed by treatment with *O*-propargylpuromycin (OPP)^48^ (5 µM, 1 h). OPP-labeled proteins were visualized by copper-catalyzed azide-alkyne cycloaddition (CuAAC) coupling to a rhodamine-azide reporter group. Results are from a single experiment representative of 3 independent experiments. **h**, Quantification of OPP signals for experiment shown in **g** and replicates. Data are average values ± SEM from 3 independent experiments and normalized to DMSO-treated parental cells. **i**, Cysteine-directed ABPP data showing enantioselective liganding of DDX49_C258 by MY-45B and MY-46B (20 µM, 3 h) in HCT116 cells. Data represent average values ± SEM of 4 independent experiments. **j**, ESCAPE results for DDX49_C258 evaluated in HCT116 cells treated with MY-45B or MY-46B (5 μM, 5 days) in HCT116. Data are average values ± SEM from 3 independent experiments. **k**, 18S rRNA signals in parental HCT116 cells treated with MY-45B or MY-46B (5-10 μM, 48 h) as measured by qPCR. Data are average values ± SEM from 4 independent experiments. All P values were calculated using two-sided Student’s t test. N.S., not significant, * p<0.05, **p<0.01, ***p<0.001.

RT-qPCR analysis revealed that MY-45B, but not MY-45A, decreased the quantity of 18S rRNA in parental HCT116 cells (**Fig. 4c** and **Supplementary Table 3**). MY-45B also suppressed 18S rRNA in WT-DDX49-expressing cells, but not in C258S-DDX49-expressing cells (**Fig. 4d**). Other rRNAs (5.8S, 28S, and 47S rRNAs) were unaffected by MY-45B (**Extended Data Fig. 4a**). We corroborated these findings with CRISPRi-mediated knockdown of endogenous DDX49, which also produced a selective depletion of 18S rRNA, but not other rRNAs in HCT116 cells (**Extended Data Fig. 4b**).

The 18S rRNA is the single RNA structural component of the 40S subunit of the ribosome^45^. Interestingly, previous unbiased genome-wide Perturb-Seq experiments have placed the gene signature associated with knocking down DDX49 within the 40S ribosomal subunit cluster^46^, supporting a potential role for DDX49 in 40S ribosome biogenesis. Also consistent with this hypothesis, genetic depletion of the yeast DDX49 ortholog Dbp8p leads to a selective deficit in 18S rRNA and the 40S ribosomal subunit^47^. Here, we found by polysome fractionation that MY-45B caused a pronounced stereoselective reduction in the 40S subunit in WT-DDX49-expressing cells, but not in C258S-DDX49-expressing cells (**Fig. 4e, f**). In contrast, MY-45B did not impair the 60S ribosomal subunit content of cells (**Fig. 4e, f**). The MY-45B-induced loss of the 40S ribosomal subunit correlated with impairments in global protein synthesis as measured by *O*-propargylpuromycin (OPP) labeling of cells^48^ (**Fig. 4g, h** and **Extended Data Fig. 4c**). As was observed for 18S rRNA and 40S ribosomal subunit, the effects of MY-45B on protein synthesis were both stereoselective and C258-dependent (**Fig. 4g, h**). These findings, taken together, indicate that MY-45B inhibits the functional contributions that DDX49 makes to 18S rRNA maturation and ribosome biogenesis, leading to impairments in both protein synthesis and cancer cell growth.

Finally, in comparing our ABPP and ESCAPE data, we noted that MY-46B, a diastereomer of MY-45B (**Fig. 3a**), also enantioselectively reacted with DDX49_C258 (**Fig. 4i**), but did not produce a resistance score indicative of impairing HCT116 cell growth (**Fig. 4j**). We confirmed that MY-46B reacted with WT-, but not C258S-DDX49 by both gel-ABPP (**Extended Data Fig. 4d**) and intact MS analysis (**Extended Data Fig. 4e**) with a similar potency to MY-45B (IC_50_ value for MY-45B: 0.76 µM; IC_50_ value for MY-46B: 1.65 µM; **Extended Data Fig. 4f**). However, MY-46B did not alter the 18S rRNA content of HCT116 cells, even when tested at a two-fold higher concentration (10 µM) than that required for MY-45B to suppress 18S rRNA (**Fig. 4k**), and MY-46B also produced only a modest reduction in cell growth that was not enantioselective and much lower in magnitude than MY-45B (**Extended Data Fig. 4g**). We interpret these results to indicate that, while both MY-45B and MY-46B enantioselectively react with C258 of DDX49, only MY-45B inhibits DDX49 activity in cells. Such divergence in the functional effects of small molecules binding at non-orthosteric sites in proteins is well-described in the literature^24–30^, and our data highlight the potential for ESCAPE to distinguish functional from silent covalent ligands engaging the same cysteine residue in cells.

## Discussion

Advanced gene editing methods, including base editing and prime editing, are increasingly being used to study how small molecules interact with proteins in human cells^49^. These studies have facilitated an understanding of the molecular basis of drug resistance^50–53^ and elucidation of noncanonical mechanisms of drug action (e.g., the inhibition of scaffolding vs catalytic functions of enzymes)^54^. ESCAPE addresses a complementary problem of assigning functions to the many covalent ligands emerging from chemical proteomics and other binder-based screening methods^24^. Such covalent ligands target proteins from diverse structural and functional classes, often at non-orthosteric sites, and ESCAPE offers a consolidated method to assay these nascent small molecule-protein interactions for impact on a measurable cellular property. Here, we have applied ESCAPE to quantify the effects of covalent ligand-protein interactions on cancer cell growth, but we envision the method being applicable to additional cellular processes, including gene regulation effects converted into positive-selection cell growth assays^55^ and cell state changes measured with sortable surface markers (e.g., immune cell activation).

Our results further highlight the potential for ESCAPE to not only identify covalent compounds acting by loss-of-function mechanisms directionally aligned with the gene dependency scores of target proteins, but also to characterize anti-proliferative agents displaying gain-of-function activities that are more challenging to predict from genetic disruption experiments (e.g., NONO ligands; **Fig. 2d, e**). We are also encouraged that, in at least one instance, ESCAPE was capable of simultaneously identifying two growth inhibitory protein interactions for the same covalent ligand (GRPEL1_C124 and SF3B1_C1111 for WX-03-58). We interpret this result to mean that ESCAPE has the potential to deconvolute the contributions of individual protein interactions to the overall cell growth effect of a multi-target compound. To the extent that this feature proves generalizable in ESCAPE assays, it should further accelerate the functional assignment of nascent liganding events by circumventing the need for extensive medicinal chemistry optimization of their proteome-wide selectivity. Nonetheless, we acknowledge that a minimal level of potency is likely required for a covalent ligand-protein interaction to be assayable by ESCAPE such that excessive electrophilic stress-related forms of toxicity are avoided. For the tryptoline acrylamides tested herein, we found that concentrations up to 10-15 µM gave interpretable resistance scores in ESCAPE assays.

Our characterization of covalent ligands targeting DDX49 supports a specific role for this RNA helicase in the maturation of 18S rRNA^47^, with the resulting impairments in 40S ribosome subunit biogenesis and protein synthesis in turn providing a mechanistic rationale for the enantioselective growth inhibitory effects of MY-45B. While we do not yet understand how MY-45B inhibits the function of DDX49, the location of C258 in the predicted protein structure may suggest that MY-45B allosterically perturbs RNA interactions. Our data further suggest that covalent ligands targeting DDX49_C258 can produce distinct functional outcomes, as the stereoisomeric azetidine butynamide MY-46B, despite engaging this site with similar potency to MY-45B, did not give a positive resistance score by ESCAPE and was not found to perturb the 18S rRNA content of cancer cells. The capacity of ESCAPE to distinguish functional from silent covalent ligands, even when they target the same cysteine, should prove particularly helpful in the characterization of non-orthosteric sites on proteins, where the structure-activity relationships for small-molecule binding and efficacy can show surprising divergence^26–29^.

We also note several limitations of ESCAPE. First, many covalent ligand-protein interactions may produce functional effects that do not impact cell growth. Thus, while restricting our ESCAPE assays to a cell proliferation readout had the advantage of narrowing the pharmacologically relevant target space for early-stage covalent ligands like the tryptoline acrylamide stereoprobes, the absence of a positive resistance score should not be interpreted as strong evidence that a given small molecule-protein interaction is non-functional. ESCAPE also depends on proteins maintaining basal functionality when ligandable cysteines are mutated to serine or alanine, and some proteins may not tolerate such mutations. Of course, the genetic essentiality of a cysteine itself would imply that covalent ligands targeting that residue have high functional potential^17,56^. Finally, ESCAPE assumes that the target engagement of electrophilic small molecules is primarily mediated by covalent binding, but some compounds could exhibit sufficient reversible binding affinity to retain activity with the Ser-edited protein. Considering that even advanced clinical-stage compounds developed by “covalent-first” ^57^ methods like ABPP lose activity when their target cysteines are mutated to serine or alanine^26,58^, we expect that ESCAPE will prove broadly applicable to covalent ligands at various stages of development.

Projecting forward, we believe that ESCAPE, when combined with the large-scale ligandability maps emerging from ABPP studies, will offer a robust chemistry-first framework for assigning functions to cryptic sites on proteins from diverse structural and mechanistic classes. As we have shown through the mapping of small molecule-protein interactions displaying both gain- and loss-of-function activities, ESCAPE has the potential to pharmacologically validate and extend beyond the outputs of global gene disruption screens. While we have focused on cysteine-directed compounds, ESCAPE should be applicable to covalent chemistry targeting additional amino acids^7,59–75^. And, recent advances in site mapping of non-covalent small molecule-protein interactions^76^ further point to the potential for adapting ESCAPE to characterize the functions of reversible ligands. Finally, integrating ESCAPE data on cryptic ligandable pockets with frontier molecular dynamics and machine learning models^77–80^ may provide a general approach for illuminating the small molecule-induced conformational transitions that underpin non-orthosteric functional sites in proteins.

## Methods

### Cell culture

Cell lines used in this study included HCT116 (#CCL-247, ATCC), KMS11 (#JCRB1179, JCRB), K562 (#CCL-243, ATCC), TC71 (Texas Tech University), OVCAR8 (DCTD Tumor Repository), NUGC3 (#JCRB0822, JCRB), CALU6 (#HTB-56, ATCC), 22RV1 (#CRL-2505, ATCC), and HEK293T (#632180, Takara). All human cell lines were authenticated by the vendors using short tandem repeat (STR) profiling. Cells were cultured in RPMI-1640 medium (Gibco) under standard conditions (37 °C, 5% CO₂) and were confirmed to be free of microbial contamination. All culture medium was supplemented with 10% fetal bovine serum (FBS; Omega Scientific), 100 U ml⁻¹ penicillin, 100 μg ml⁻¹ streptomycin (Gibco), and 2 mM GlutaMAX (Gibco).

### Cloning of pegRNA

PLV107 vector (20 μg) was digested with BsmBI (20 μl; NEB) in a total reaction volume of 200 μl using NEBuffer 3.1 at 37 °C for 2 hours. The linearized vector was resolved by 1% agarose gel electrophoresis and purified using the QIAquick Gel Extraction Kit (Qiagen). The pegRNA sequences were designed according to established prime editing principles^81^. Gene fragments encoding pegRNAs were cloned into PLV107 by Golden Gate assembly. Briefly, 50 ng of linearized PLV107 was combined with 5 ng insert DNA (1 μl), 1 μl of 10 mM DTT, 1 μl of 10 mM ATP, 0.2 μl Esp3I (Thermo Scientific), and 0.2 μl T7 DNA ligase (Qiagen Beverly) in 1 μl of 10× Tango buffer, in a total volume of 10 μl. The reaction was repeated between 37 °C (3 min) and 20 °C (3 min) for 30 cycles, followed by incubation at 37 °C for 30 min and 65 °C for 10 min to inactivate the enzymes. The ligation mixture was then used directly for transformation into competent stbl3 cells.

### Lentivirus particle production

Plasmids, including the transfer vector, lentiviral packaging vector (pCMV-dR8.91), and envelope vector (VSV-G), were mixed at a 6:6:1 mass ratio in Opti-MEM (Gibco). Polyethylenimine (PEI; Polysciences) was added at 3 μl of 1 μg μl⁻¹ stock per μg of total DNA. After incubation at room temperature for 20 min, the transfection mixture was added dropwise to HEK293T cells (Takara) at ∼50% confluence. After 8 h, the medium was replaced with fresh DMEM (Corning) supplemented with 30% fetal bovine serum (FBS), 100 U ml⁻¹ penicillin, 100 μg ml⁻¹ streptomycin, and 2 mM GlutaMAX (Gibco). Viral supernatants were collected 48 h post-transfection and clarified by filtration through 0.45 μm syringe filters (Millipore) to remove cellular debris.

### ESCAPE assay

Cells with stable integration of NG-PAM compatible PE7^36,38,82^ using lentivirus were seeded on day 0 in 96-well plates at a density of 5,000–20,000 cells per well and transduced with pegRNA^35^ viral supernatant supplemented with 8 μg/mL polybrene (Millipore). After allowing cells to settle, spin infection was performed at 900 × g for 1 h at 30 °C. On day 1, virus-containing medium was replaced with fresh culture medium. After 7 days, transduced cells were re-plated into new 96-well plates at a density of 5,000–20,000 cells per well (adjusted based on growth rate) prior to ligand treatment. Cells were treated for another 5 days in a tissue culture incubator (37 °C, 5% CO₂) before collection.

For genomic DNA extraction, ligand-containing medium was removed, followed by a brief Phosphate-Buffered Saline (PBS) wash. Cells were lysed in 40–100 μL lysis buffer (10 mM Tris-HCl, pH 7.5, 0.5% Tween-20, 0.02% SDS) supplemented with freshly added proteinase K (20 μg/mL). Plates were incubated at 55 °C for 10 min, and lysates were transferred to a 96-well PCR plate and sealed with adhesive foil. Samples were further digested at 55 °C for 1.5 h, followed by heat inactivation at 95 °C for 30 min. Plates were briefly centrifuged to collect condensation.

Genomic regions containing edited sites were amplified and indexed using a two-step PCR protocol. For PCR1, 5 μL genomic DNA lysate, 12.5 μL Phusion PCR Master Mix (Thermo Fisher), and 1.25 μL each of 10 μM forward and reverse primers (containing Illumina adapter sequences) were combined to a final volume of 25 μL. PCR1 conditions were: 95 °C for 3 min; 30 cycles of 95 °C for 20 s, 60 °C for 25 s, and 72 °C for 25 s; followed by a final extension at 72 °C for 2 min. PCR1 products were purified using Mag-Bind® TotalPure beads (Omega Bio-tek) according to the manufacturer’s instructions using multichannel pipettors and eluted in 50 μL of 10 mM Tris-HCl (pH 7.5).

For PCR2, 5 μL purified PCR1 product, 12.5 μL Phusion PCR Master Mix, and 1.25 μL each of 10 μM forward and reverse indexing primers were combined in a 25 μL reaction. PCR2 conditions were: 98 °C for 3 min; 12 cycles of 95 °C for 20 s, 60 °C for 25 s, and 72 °C for 25 s; followed by a final extension at 72 °C for 2 min. PCR2 products were pooled and purified using Mag-Bind® TotalPure beads.

Libraries were quantified using Qubit assay kits (Invitrogen) and sequenced on an Illumina MiniSeq platform (GenerateFASTQ v2.0.1) with a 10% PhiX spike-in. Paired-end reads were demultiplexed based on combinatorial dual indices. Genome editing outcomes were quantified using CRISPResso2^83^ with the following parameters: CRISPResso --fastq_r1 {r1_file} --fastq_r2 {r2_file} --amplicon_seq {amplicon_WT} --expected_hdr_amplicon_seq {amplicon_edit} –g {guide} -wc -3 -w 20 -q 30 --discard_indel_reads --min_paired_end_reads_overlap 1

### Generation of clonal C1111S-SF3B1 HCT116 cells

HCT116 cells stably expressing NG-PAM compatible PE7^36,38,82^ and a pegRNA^35^ creating C1111S-SF3B1 (**Supplementary Table 1**) were generated as described above. Single cells were isolated by fluorescence-activated cell sorting using a MoFlo Astrios EQ jet-in-air sorter into individual wells of three 96-well plates containing RPMI media. Single-cell clones were expanded and genotyped by targeted genomic PCR as described in the ESCAPE method section above. Validated five homozygous C1111S-SF3B1 clones were then pooled and used for the sensitivity evaluation assay as described in **Fig. 1**.

### Western blot analysis

Cells were collected by centrifugation (500 × g, 5 min) and washed with PBS. Proteins were extracted using RIPA lysis buffer and quantified using a BCA assay kit (Thermo Fisher Scientific). Lysates were mixed with 4× LDS sample buffer (Invitrogen), resolved by SDS–PAGE, and transferred onto PVDF membranes using a Mini Trans-Blot® Cell system (Bio-Rad). Blots were blocked in TBST buffer (20 mM Tris-HCl, pH 7.5, 150 mM NaCl, 0.1% Tween-20) containing 5% non-fat milk for 1 h at room temperature. Blots were incubated overnight at 4 °C with primary antibodies, including V5-HRP (A190-120P, Bethyl Laboratories), GAPDH-HRP (HRP-60004, Proteintech), and DDX49 (ab188010, Abcam), followed by three washes with TBST (5 min each). For DDX49, the blot was then incubated with anti-rabbit-HRP (SA00001-2, Proteintech). After three additional TBST washes (5 min each), signals were developed using chemiluminescent HRP substrates (Thermo Fisher Scientific) and imaged using a Bio-Rad ChemiDoc imaging system.

### Gel-ABPP

Approximately 2 million HCT116 cells stably expressing V5-tagged DDX49 were treated with competitor probes followed by alkyne probes, or with alkyne probes alone, in RPMI media. After treatment, cells were collected, washed with ice-cold PBS, and lysed in 300 μl of lysis buffer (25 mM Tris-HCl, pH 7.5, 150 mM NaCl, 1% Triton X-100) followed by probe sonication (20 pulses, 10% power output). Lysates were clarified by centrifugation at 16,000 × g for 5 min, and soluble protein fractions were normalized. Ten percent of the normalized lysate was reserved as input, and the remainder was incubated with anti-V5 magnetic beads (Proteintech) at 4 °C for 2 h. Beads were washed twice with wash buffer (25 mM Tris-HCl, pH 7.5, 150 mM NaCl, 0.1% Triton X-100). For click chemistry labeling, a 5.5 μl reaction mix containing 1 μl CuSO_4_ (50 mM stock), 3 μl TBTA ligand (1.7 mM stock in DMSO:tBuOH, 1:4 v/v), 1 μl freshly prepared TCEP (50 mM stock), and 0.5 μl rhodamine-azide (1.25 mM stock) was mixed separately in order and then added to each sample in 50 μl PBS. After 1 h, samples were mixed with NuPAGE LDS sample buffer, heated at 90 °C for 5 min, and analyzed by SDS–PAGE. Rhodamine fluorescence signal was acquired using a ChemiDoc MP imaging system (Bio-Rad).

### MS-ABPP

Approximately 1-2 × 10^7^ cells were treated with the indicated compounds for the specified concentrations and durations. Cells were harvested by scraping or centrifuging, washed with ice-cold PBS, and stored at −80 °C prior to further processing. Detailed protocols for protein-directed and cysteine-directed ABPP, as well as mass spectrometry instrumentation parameters, have been described previously^9,11,16,32^.

### Recombinant DDX49 purification

HEK293T cells were seeded in antibiotic-free DMEM. Cells at ∼40% confluence were transfected with pTwist_CMV_WPRE_Neo plasmids encoding WT-DDX49 or C258S-DDX49 using the TransIT-X2 Dynamic Delivery System (Mirus) according to the manufacturer’s instructions. Three days post-transfection, cells were treated with 10 μM MY-45A, MY-45B or DMSO for 3 hours, washed twice with PBS (600 × g, 5 min each), and stored at −80 °C until further use.

The base buffer for DDX49 purification consisted of 25 mM HEPES (pH 7.5), 150 mM NaCl, and 0.1 mM DTT. Frozen cell pellets were lysed in base buffer supplemented with 1% Triton X-100 using a probe sonicator (1 s on, 1 s off; total 4 min on-time; 20% amplitude). Lysates were first clarified by centrifugation at 3,000 × g for 5 min at 4 °C, followed by high-speed centrifugation at 30,000 × g for 20 min at 4 °C. Twin-Strep-tagged DDX49 proteins were purified using Strep-Tactin® 4Flow® high-capacity resin (IBA) according to the manufacturer’s protocol. After incubation with the resin for 3 hours at 4 °C with gentle agitation, beads were collected using Econo-Pac® columns (Bio-Rad) and washed extensively with base buffer containing 0.1% Triton X-100. Bound proteins were eluted twice with base buffer supplemented with 10 mM desthiobiotin. Eluted proteins were concentrated using PES protein concentrators (Pierce), and concentrations were determined using a 660 nm protein assay (Pierce). Samples were aliquoted and stored at −80 °C until use. The protein samples were desalted using Zeba spin columns (7K MWCO; ThermoFisher, 89883) according to the manufacturer’s protocol prior to intact MS analysis, as described previously^32^.

### rRNA qPCR

HCT116 cells were seeded at a density of 3.0 × 10^5^ cells per well in 6-well plates and treated with ligands at the time of plating. After 24 h, media and compounds were refreshed. Following a total of 48 h of treatment, cells were harvested in TRIzol (Thermo Fisher Scientific) for RNA isolation. Total RNA was purified using the Direct-zol™ RNA MiniPrep Plus Kit (Zymo Research) according to the manufacturer’s instructions.

For reverse transcription, 500 ng of total RNA per reaction was used. RNA was combined with random hexamers (1 μL, 20×), dNTPs (1 μL, 10 mM each), and nuclease-free water to a final volume of 15 μL, incubated at 65 °C for 5 min, and immediately placed on ice. Reverse transcription was performed by adding 4 μL of 5× RT buffer, 0.5 μL murine RNase inhibitor (New England Biolabs; diluted 1:5), and 0.5 μL Maxima H Minus Reverse Transcriptase (Thermo Fisher). Reactions were incubated at 25 °C for 10 min, followed by 50 °C for 30 min, and heat-inactivated at 85 °C for 5 min. The resulting cDNA was diluted 1:5 with nuclease-free water (20 μL RT product + 80 μL H_2_O) and stored at −20 °C until use.

Quantitative PCR (qPCR) was performed using Power SYBR Green PCR Master Mix (Thermo Fisher) on a QuantStudio 5 Real-Time PCR System (Thermo Fisher) in 384-well plates. Each 10 μL reaction contained 5 μL of 2× Power SYBR Green Master Mix, 3.6 μL nuclease-free water, and 0.4 μL primer mix (final concentration 400 nM for each primer). One microliter of 1:5 diluted cDNA was added per reaction. All samples were analyzed in technical duplicates with at least three biological replicates per condition. Cycling conditions were 95 °C for 10 min, followed by 30 cycles of 95 °C for 15 s and 60 °C for 1 min. Relative gene expression was calculated using the ΔΔCt method, normalized to GAPDH, and expressed relative to the DMSO control.

### Protein synthesis OPP labeling assay

HCT116 cells were seeded at a density of 3.0 × 10^5^ cells per well in 6-well plates and treated with DMSO, 5 μM MY-45A, or 5 μM MY-45B. After 24 h, media and compounds were refreshed, and cells were cultured for a total of 48 h. Where indicated, cells were treated with cycloheximide (50 μg/mL) for 15 min at 37 °C as a control for protein synthesis inhibition. Cells were then incubated with 5 μM *O*-propargylpuromycin (OPP) for 1 h at 37 °C. Following OPP labeling, cells were harvested, washed with PBS, and pellets were stored at −80 °C. Cell pellets were lysed in lysis buffer (25 mM Tris-HCl, pH 7.4, 150 mM NaCl, 1% Triton X-100, 10% glycerol, 1 mM MgCl₂) supplemented with protease inhibitor cocktail and benzonase. Lysates were sonicated, followed by addition of SDS to a final concentration of 0.5%, gentle vortexing, and incubation at 4 °C for 5 min. Samples were clarified by centrifugation at 20,000 × g for 5 min at 4 °C. Protein concentrations were determined by BCA assay, and samples were normalized to 1.0 μg/μL. Equal amounts of protein were subjected to copper-catalyzed azide–alkyne cycloaddition (CuAAC) to conjugate OPP-labeled nascent polypeptides with rhodamine azide (Rh-N₃). A click reaction master mix was prepared immediately prior to use by sequential addition of CuSO₄ (1 μL of 50 mM stock), Tris[(1-benzyl-1H-1,2,3-triazol-4-yl)methyl]amine (TBTA; 3 μL of 1.7 mM solution in DMSO:tBuOH, 1:4, v/v), tris(2-carboxyethyl)phosphine (TCEP; 1 μL of freshly prepared 50 mM solution), and rhodamine azide (0.4 μL of 1.25 mM stock), with vortexing and brief centrifugation after each addition. For each sample, 5 μL of the master mix was added to 50 μL of lysate, followed by gentle vortexing and incubation at room temperature for 1 h. Reactions were terminated by addition of NuPAGE LDS sample buffer (Invitrogen) containing β-mercaptoethanol. Samples were resolved by SDS–PAGE and imaged for rhodamine fluorescence to detect OPP incorporation. Gels were subsequently stained with Coomassie Brilliant Blue overnight and destained overnight prior to imaging.

### Polysome fractionation

HCT116 cells were seeded at a density of 2.0 × 10^5^ cells/mL in 15-cm dishes and treated with DMSO, 5 μM MY-45A, or 5 μM MY-45B at the time of plating. After 24 h, media and compounds were refreshed, and cells were harvested following a total of 48 h of treatment. Prior to harvest, cells were treated with cycloheximide (100 μg/mL) for 5 min at 37 °C. Cells were then washed twice with ice-cold PBS supplemented with cycloheximide (100 μg/mL), scraped, and pelleted by centrifugation (300 × g, 5 min, 4 °C). Cell pellets were snap-frozen in liquid nitrogen and stored at −80 °C.

Sucrose gradients (10–50%) were prepared using a Gradient Master in polysome buffer containing 20 mM Tris-HCl (pH 7.5), 150 mM NaCl, 5 mM MgCl₂, 1 mM DTT, 100 μg/mL cycloheximide, protease inhibitor cocktail, and 20 U/mL murine RNase inhibitor. Frozen cell pellets were thawed on ice and lysed in 200 μL of polysome buffer supplemented with 0.5% NP-40. Lysates were clarified by centrifugation (20,000 × g, 5 min, 4 °C) and normalized to total RNA concentration within each experiment using a NanoDrop spectrophotometer. Equal amounts of RNA were loaded per condition within each replicate (typically ∼200–240 μg RNA per gradient) and 200 μL of lysate was loaded to each sucrose gradient. Gradients were subjected to ultracentrifugation in an SW41 Ti rotor (35,000 rpm, 4 °C, 3 h). Following centrifugation, gradients were fractionated using a Biocomp Gradient Station, and absorbance at 260 nm was continuously monitored using a Triax flow cell.

### Genome-wide CRISPRi screens

The human CRISPRi sgRNA library Dolcetto^41^ cloned into the pXPR_050 backbone was used for pooled screening. On day 0, 1.2 × 10^8^ HCT116-pZR071-dCas9-KRAB cells, with or without recombinant DDX49 expression, were seeded into 12-well plates at a density of 2 × 10^6^ cells per well and transduced with Dolcetto library viral supernatant supplemented with 8 μg/mL polybrene. Spin infection was performed at 900 × g for 2 h. At 16 h post-infection, cells were transferred to 15-cm dishes. On day 2, puromycin selection (1.7 μg/mL) was initiated to enrich for sgRNA-transduced cells; the infection rate was maintained at ∼50% to achieve an optimal multiplicity of infection. On day 5, cells were split into treatment arms and treated with 5 μM MY-45A, MY-45B, or DMSO control. Cells were cultured for an additional 12 days with media changes every 3 days, including fresh compound. Genomic DNA was isolated from cell pellets using NucleoSpin Blood L kits (Macherey-Nagel) according to the manufacturer’s instructions and quantified using the PicoGreen dsDNA Assay Kit (Thermo Fisher Scientific) before genomic PCR^17^.

### Statistics and Reproducibility

Statistical analyses in this paper were performed using R (v4.5.1). Data visualization was done in R using package *tidyplots*. To compare the means between two groups of data points, two-sided Student’s t test was used to calculate the P values. Two-sided statistical tests were performed unless stated otherwise. The Benjamini–Hochberg procedure was used to adjust multiple hypothesis testing when applicable.

## Supporting information

Supplementary Chemistry

## Acknowledgements

We thank WuXi AppTec for the synthesis of chemical probes used in this study and the Scripps Automated Synthesis Facility for HRMS measurements. We thank Dr. Luke Wiseman for sharing his qPCR machine and Dr. Luke Lairson for sharing his Envision plate reader. We acknowledge this work was supported by the Damon Runyon Cancer Research Foundation (DRG: 2406-20 awarded to H.L.), NCI (R35 CA231991 awarded to B.F.C and K99/R00 CA290143 awarded to H.L.). NIGMS (R35 GM136412 awarded to J.R.W.). D.R.L. is supported by NIH R35 GM118062, RM1 HG009490, and HHMI.

## Author Contributions Statement

Conceptualization: B.F.C., H.L.; Methodology: J.E.T., B.M., D.R.L., B.F.C, H.L.; Investigation: J.E.T., B.M., W.R.B., R.E.H., K.R.S., K.E.D, E.N., K.S., J.R.W., H.L.; Visualization: J.E.T., B.M., H.L.; Funding acquisition: B.F.C., H.L., D.R.L., G.W.Y.; Project administration: S.B.; Supervision: B.F.C., H.L., D.R.L., G.W.Y.; Writing: B.F.C., H.L.

## Data availability

Proteomics data have been deposited to the ProteomeXchange Consortium (PXD081738). Sequencing data have been deposited in the NCBI Sequence Read Archive (PRJNA1501859). Processed screen data and proteomics data are provided as **Supplementary Table 2**. Source data are provided with this paper. For cancer dependency chronos scores, DepMap (https://depmap.org/portal/, 25Q2) was used.

## Code availability

Custom code used in the analysis is available on github (https://github.com/jasonli1314/ESCAPE).

---

**Extended Data Fig. 1.**
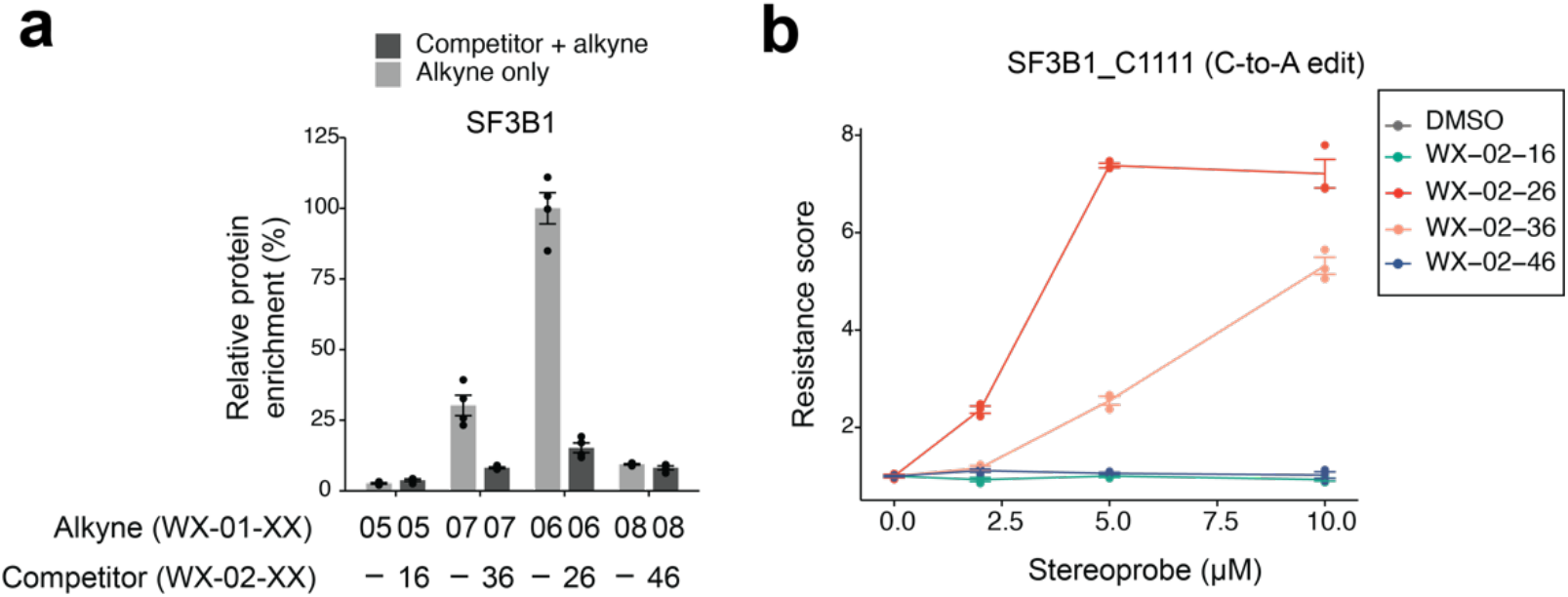
Benchmarking of ESCAPE. **a**, Protein-directed ABPP data derived from ref.^11^ showing enantioselective enrichment of SF3B1 in Ramos cells by alkyne stereoprobes WX-01-06 and WX-01-07 (5 µM, 1 h) and blockade of this enrichment by the corresponding parent stereoprobes WX-02-26 and WX-02-36 (20 µM, 2 h pre-treatment). See Fig. 1b for structures of stereoprobes. Data are average values ± SEM from 4 independent experiments. **b**, ESCAPE assay performed in HCT116 cells harboring the C1111A-SF3B1 edit treated with the tryptoline acrylamide stereoprobes from Fig. 1b at the indicated concentrations. Data are average values ± SEM from 3 independent experiments.

**Extended Data Fig. 2.**
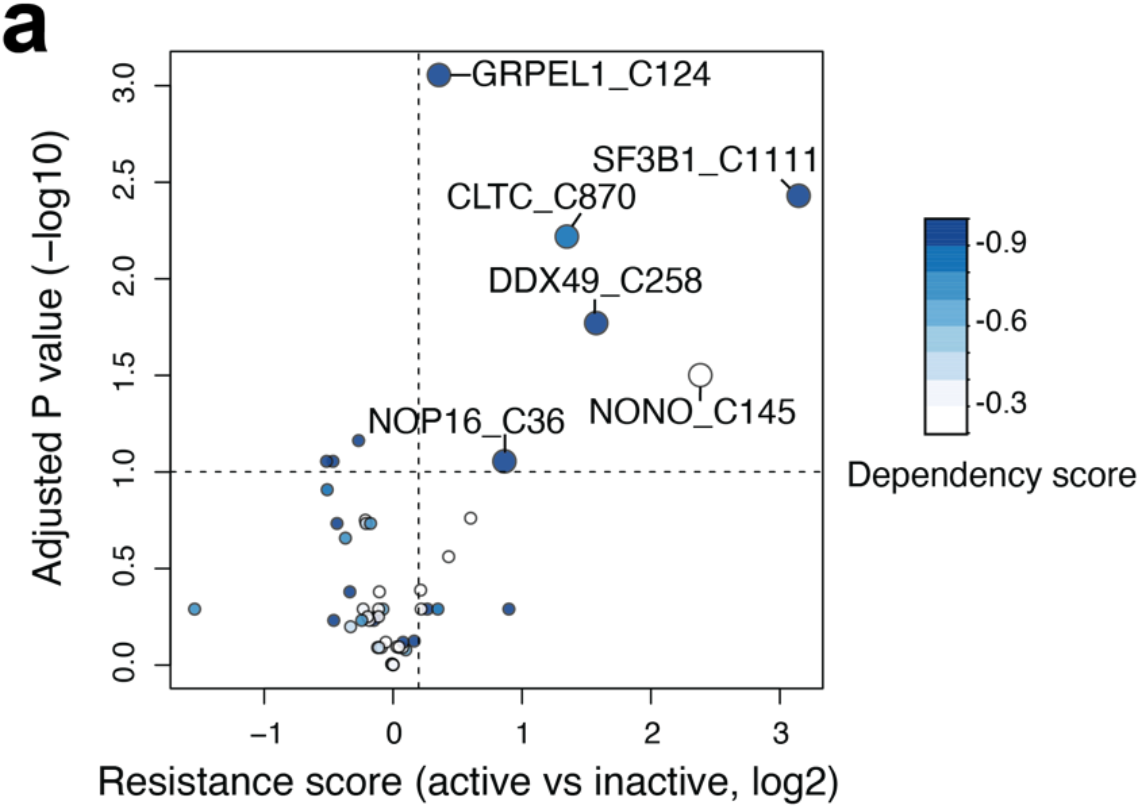
ESCAPE identifies covalent liganding events that impair cancer cell growth. **a**, Volcano plot showing the resistance scores and adjusted P values for the covalent ligand-protein interactions assayed by ESCAPE in HCT116 cells treated with active vs inactive stereoprobes. Prime-edited HCT116 cells harboring Cys-to-Ser point mutations for each protein target were treated with respective active stereoprobe ligands or their inactive controls (5-15 µM, see **Table 1**) or DMSO for 5 days, followed by targeted genomic PCR (see **Supplementary Table 1** and **Methods** section). Dots for covalent ligand-protein interactions defined as hits in the ESCAPE assay (resistance scores (log2) > 0.2 with adjusted P values (-log10) > 1) are enlarged for visualization purposes.

**Extended Data Fig. 3.**
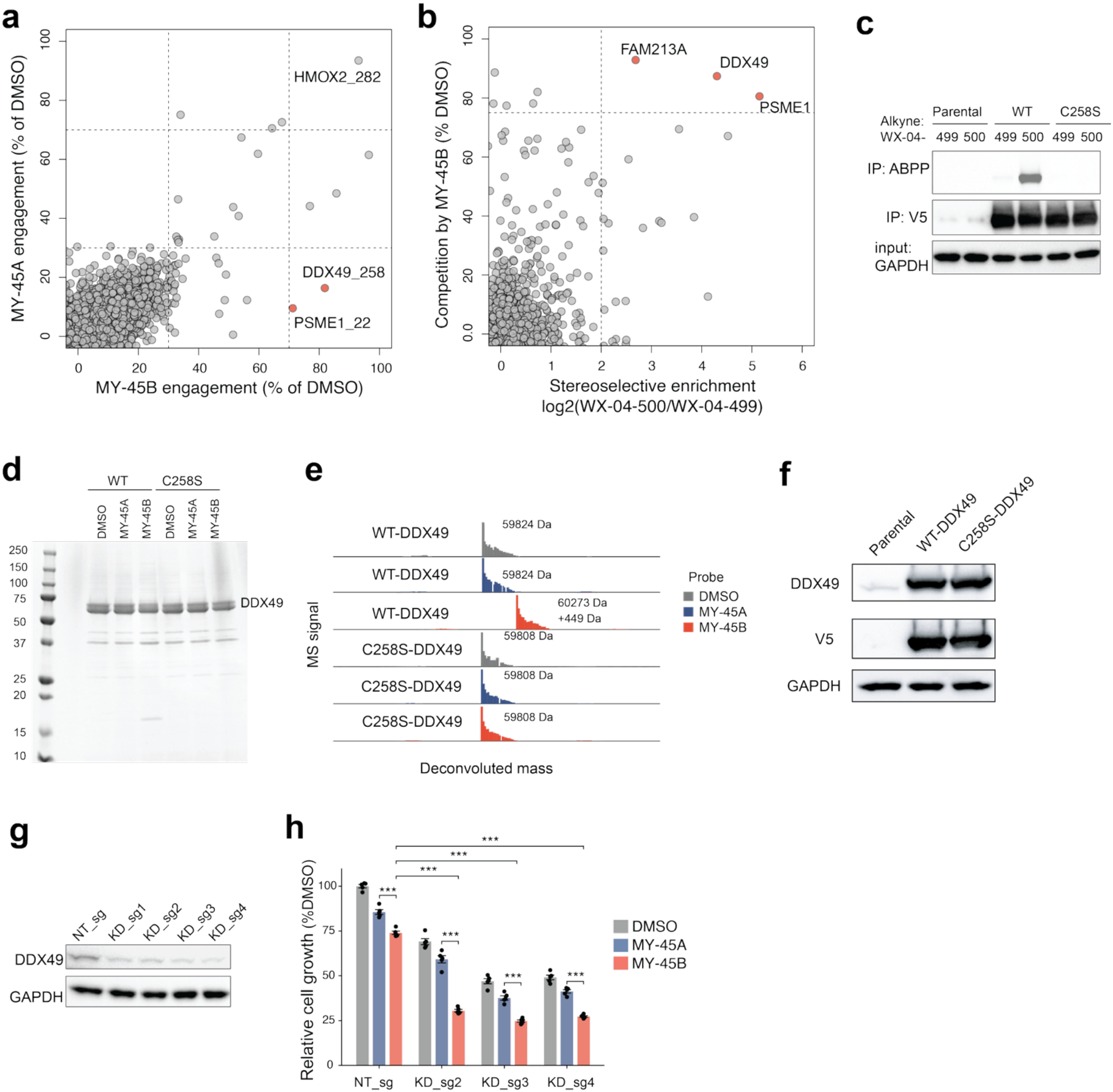
Stereoprobes targeting DDX49 perturb cancer cell growth. **a**, Scatter plot showing cysteine-directed ABPP data from ref.^16^ corresponding to 22Rv1 cells treated with MY-45B (x-axis) or MY-45A (y-axis) (5 μM, 3 h) or DMSO. Data are average values from 4 independent experiments. **b**, Protein-directed ABPP data comparing the stereoselective enrichment of proteins by WX-04-500 vs WX-04-499 (5 μM, 1 h) from HCT116 cells and the blockade of WX-04-500 enrichment of proteins by pre-treatment with MY-45B (5 μM, 2 h). Data are average values from 2 independent experiments. **c**, Gel-ABPP data showing enantioselective engagement of recombinant WT-DDX49, but not C258S-DDX49, by alkyne stereoprobe WX-04-500. Parental, V5-tagged WT-DDX49-expressing, or V5-tagged C258S-DDX49-expressing HCT116 cells were treated with WX-04-500 or enantiomer WX-04-499 (2 μM, 1 h), lysed, and V5-tagged DDX49 proteins were immunoprecipitated with an anti-V5 antibody prior to analysis by gel-ABPP. Data are from a single experiment representative of 2 independent experiments. **d**, Purified human DDX49 analyzed by SDS-PAGE and Coomassie blue staining. Twin-strep-tagged DDX49 variants were recombinantly expressed in HEK293T cells, treated with the indicated stereoprobes (10 μM, 3 h *in cellulo*) or DMSO control. Then cells were lysed and DDX49 proteins were purified as described in the **Methods**. **e**, Intact protein MS data for purified stereoprobe-treated WT-DDX49 and C258S-DDX49 proteins shown in panel **d**. Proteins were analyzed by time-of-flight (TOF)-LC/MS, and data shown are deconvoluted mass spectra from a single experiment. **f**, Western blot showing recombinant WT- and C258S-DDX49 expression in HCT116 cells as measured with anti-DDX49 (upper blot) or anti-V5 (middle blot) antibodies. Results are from a single experiment representative of 2 independent experiments. **g**, Western blot showing DDX49 expression in HCT116-dCas9-KRAB cells treated with non-targeting (NT) or DDX49-targeting CRISPRi knockdown (KD) sgRNAs for 6 days. **h**, Relative growth of HCT116-dCas9-KRAB cells treated with non-targeting (NT) or DDX49-targeting CRISPRi sgRNAs (6 days) followed by treatment with a low concentration (2 µM, 4 days) of MY-45B or MY-45A. Growth effects were measured by CellTiter-Glo and shown relative to DMSO-treated NT_sg-expressing cells. Data are average values ± SEM from 5 independent experiments. All P values were calculated using two-sided Student’s t test. N.S., not significant, * p<0.05, **p<0.01, ***p<0.001.

**Extended Data Fig. 4.**
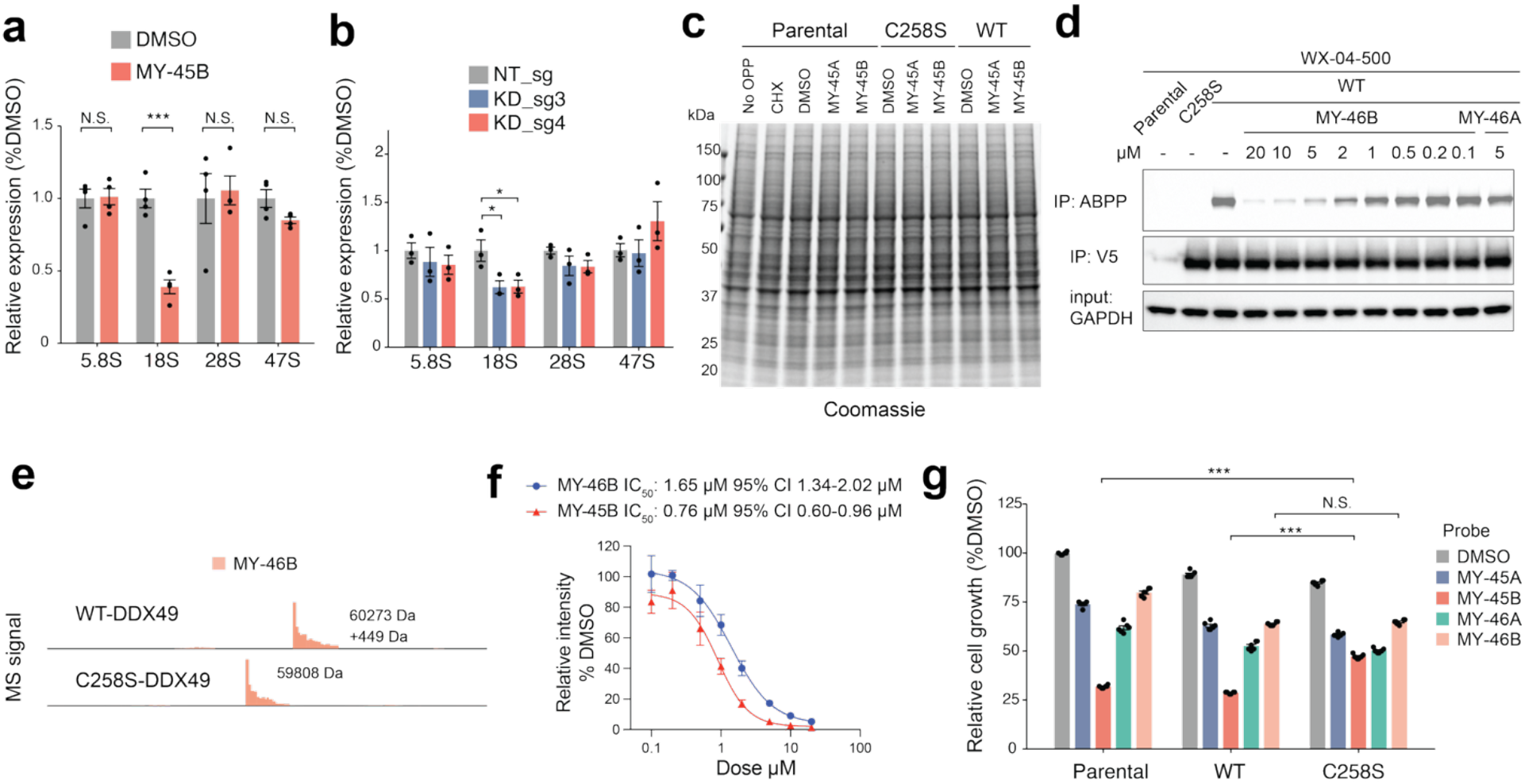
MY-45B impairs ribosome biogenesis and protein synthesis in cancer cells. **a**, Signals for the indicated rRNAs in parental HCT116 cells treated with MY-45B (5 μM, 48 h) as measured by qPCR. Data are average values ± SEM from 4 independent experiments. **b**, Effect of CRISPRi-mediated DDX49 knockdown on the indicated rRNA signals in HCT116-dCas9-KRAB cells as measured by qPCR (8 days after spin-infection). Data are average values ± SEM from 3 independent experiments. **c**, Coomassie blue stain of SDS-PAGE gel corresponding to the OPP analysis shown in Fig. 4g. **d**, Gel-ABPP data showing engagement of recombinant WT-DDX49, but not a C258S-DDX49 mutant, by WX-04-500 and the concentration-dependent blockade of the WX-04-500-WT-DDX49 interaction by MY-46B, but not enantiomer MY-46A. Parental, WT-DDX49-, and C258S-DDX49-expressing HCT116 cells were treated with MY-46A or MY-46B (3 h), followed by alkyne WX-04-500 (2 μM, 1 h) and analysis by gel-ABPP. Data are from a single experiment representative of 2 independent experiments. **e**, Intact protein MS data for MY-46B-treated WT-DDX49 and C258S-DDX49 proteins purified as described in **Methods**. Proteins were analyzed by time-of-flight (TOF)-LC/MS, and data show deconvoluted mass spectra from a single experiment. **f**, IC_50_ curves reflecting quantification of gel-ABPP data for WT-DDX49-expressing HCT116 cells treated with the indicated concentrations of MY-45B and MY-46B. Representative gel-ABPP data are shown in Fig. 3d (MY-45B) and **Extended Data Fig. 4d** (MY-46B). Data are normalized to DMSO-treated WT-DDX49-expressing HCT116 cells and represent average values ± SEM for 2 independent experiments. **g**, Relative growth effects of parental, WT-DDX49-expressing, or C258S-DDX49-expressing HCT116 cells treated with the indicated stereoprobes (5 µM, 4 days). Growth effects were measured by CellTiter-Glo. Data are average values ± SEM from 6 independent experiments and shown relative to parental cells treated with DMSO. All P values were calculated using two-sided Student’s t test. N.S., not significant, * p<0.05, **p<0.01, ***p<0.001.

