## Supplementary Chemistry for "ESCAPE: assigning site-specific activity to covalent ligands in cells by prime editing"

### Supplementary Information: Synthesis and Characterization of Novel Compounds

#### General considerations

All NMR spectra were recorded at 298 K unless otherwise noted. $^{1}H$ NMR spectra were recorded on Bruker Avance III 400, Avance III HD 400, Avance Neo 400 spectrometers or a ZKNJ QOne Quantum-I Plus 400 spectrometer ($^{1}H$, 400 MHz). $^{1}H$ NMR data are reported as follows: chemical shift (δ), multiplicity (s = singlet, d = doublet, t = triplet, m = multiplet; br. = broad), coupling constants, and integration. Chemical shifts are reported in parts per million (ppm) using the appropriate solvent as reference.[1] Analytical supercritical fluid chromatography (SFC) was performed on a Shimadzu LC system (flow rate: 3 mL/min, back pressure: 100 Bar, column temperature: 35 °C) equipped with a polydiode array detector. Tandem liquid chromatography/mass spectrometry (LC-MS) was performed on a Shimadzu LC-20AD series LC system equipped with a SPD-M20A polydiode array and an LCMS-2020 mass detector. Mass measurements for high-resolution mass spectrometry (HRMS) were performed on a Waters Xevo G2-XS TOF calibrated against sodium formate clusters and using a LeuEnk lockmass. Expected monoisotopic masses were calculated using MassLynx 4.1 and the m/z values for calibrant and lockmass were MassLynx-default values.

#### Experimental procedures and analytical data

The compounds used in this study were synthesized by adapting previously reported protocols.[2-5] Experimental procedures as well as analytical data for all new stereoprobes are provided.

*Synthesis of MY-46A and MY-46B*

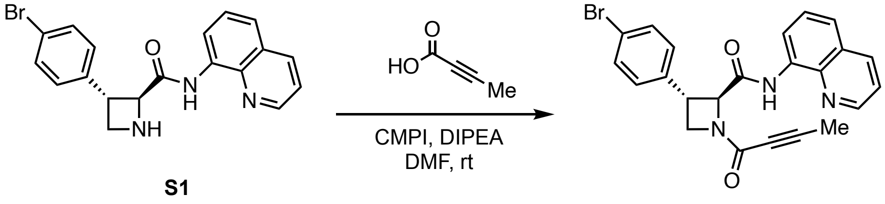


**MY-46A**

(2*S*,3*S*)-3-(4-bromophenyl)-1-(but-2-ynoyl)-*N*-(quinolin-8-yl)azetidine-2-carboxamide (MY-46A)
To a solution of S1 (50.0 mg, 131 µmol, 1 equiv), DIPEA (50.7 mg, 392 µmol, 3 equiv), but-2-ynoic acid (13.2 mg, 157 µmol, 1.2 equiv) in DMF (2 mL) was added CMPI (66.8 mg, 262 µmol, 2 equiv). The mixture was stirred at rt for 1 h. Upon reaction completion, the mixture was concentrated under reduced pressure. The resulting residue was purified by prep-HPLC (mobile phase: [A: water (0.05% $NH_4OH$), B: acetonitrile]; gradient elution: 45%-70% B over 9 min) to obtain MY-46A (16.0 mg, 27% yield) as a white solid.
$^{1}H$ NMR (400 MHz, $CD_3OD$): δ 8.86 (dd, *J* = 4.0, 1.9 Hz, 1H), 8.72 (t, *J* = 8.7 Hz, 1H), 8.38 – 8.26 (m, 1H), 7.72 – 7.50 (m, 5H), 7.40 (d, *J* = 8.5 Hz, 2H), 5.29 (d, *J* = 5.7 Hz, 0.45H), 5.10 (d, *J* = 6.3 Hz, 0.55H), 4.65 (t, *J* = 8.9 Hz, 0.55H), 4.52 (t, *J* = 9.4 Hz, 0.45H), 4.34 (dd, *J* = 8.8, 6.5 Hz,

0.55H), 4.18 – 4.08 (m, 1H), 4.07 – 3.97 (m, 0.45H), 2.10 (s, 1.65H), 1.73 (s, 1.35H); 1 exchangeable proton not observed; mixture of rotamers.
HRMS m/z calc. for $C_{23}H_{19}BrN_3O_2$ $[M+H]^+$ 448.0661 found 448.0670.

(2*R*,3*R*)-3-(4-bromophenyl)-1-(but-2-ynoyl)-*N*-(quinolin-8-yl)azetidine-2-carboxamide (MY-46B)
Prepared in similar fashion from *ent*-S1.
$^1$H NMR (400 MHz, $CD_3OD$): δ 8.87 (dd, *J* = 4.3, 1.6 Hz, 1H), 8.76 – 8.67 (m, 1H), 8.32 (dd, *J* = 10.7, 7.9 Hz, 1H), 7.73 – 7.49 (m, 5H), 7.40 (d, *J* = 8.3 Hz, 2H), 5.28 (d, *J* = 5.7 Hz, 0.45H), 5.10 (d, *J* = 6.3 Hz, 0.55H), 4.65 (t, *J* = 8.9 Hz, 0.55H), 4.52 (t, *J* = 9.4 Hz, 0.45H), 4.38 – 4.30 (m, 0.55H), 4.18 – 4.09 (m, 1H), 4.07 – 3.98 (m, 0.45H), 2.10 (s, 1.65H), 1.73 (s, 1.35H); 1 exchangeable proton not observed; mixture of rotamers.
HRMS m/z calc. for $C_{23}H_{19}BrN_3O_2$ $[M+H]^+$ 448.0661 found 448.0663.

*Synthesis of WX-04-499 and WX-04-500*

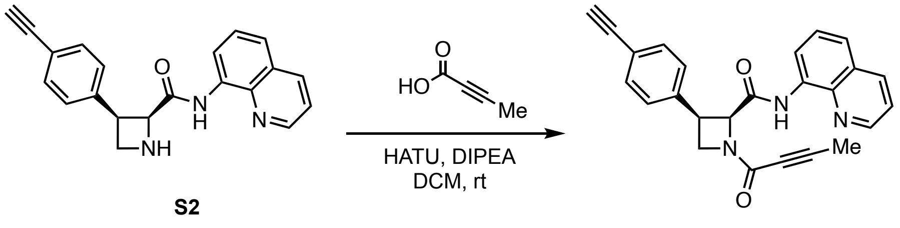


(2*S*,3*R*)-1-(but-2-ynoyl)-3-(4-ethynylphenyl)-*N*-(quinolin-8-yl)azetidine-2-carboxamide (WX-04-499)
To a solution of S2 (45.0 mg, 137 µmol, 1 equiv) and but-2-ynoic acid (17.3 mg, 206 µmol, 1.5 equiv) in DCM (2 mL) were added DIPEA (53.3 mg, 412 µmol, 3 equiv) and HATU (78.4 mg, 206 µmol, 1.5 equiv). The mixture was stirred at rt for 2 h. Upon reaction completion, the mixture was concentrated under reduced pressure. The resulting residue was purified by prep-HPLC (mobile phase: [A: water (0.05% $NH_4OH$), B: acetonitrile]; gradient elution: 36%-66% B over 9 min) to give WX-04-499 (26.4 mg, 49% yield) as a white solid.
$^1$H NMR (400 MHz, $CD_3OD$): δ 8.90 – 8.83 (m, 1H), 8.27 (t, *J* = 6.7 Hz, 1H), 8.20 (dd, *J* = 15.0, 7.7 Hz, 1H), 7.63 – 7.50 (m, 2H), 7.43 (q, *J* = 7.6 Hz, 1H), 7.36 (dd, *J* = 8.3, 3.9 Hz, 2H), 7.19 (d, *J* = 8.0 Hz, 2H), 5.62 (d, *J* = 7.9 Hz, 0.55H), 5.40 (d, *J* = 9.7 Hz, 0.45H), 4.71 – 4.57 (m, 1H), 4.49 – 4.36 (m, 2H), 3.34 (s, 1H), 2.13 (s, 1.35H), 1.83 (s, 1.65H); 1 exchangeable proton not observed; mixture of rotamers.
HRMS m/z calc. for $C_{25}H_{20}N_3O_2$ $[M+H]^+$ 394.1556 found 394.1558.

(2*R*,3*S*)-1-(but-2-ynoyl)-3-(4-ethynylphenyl)-*N*-(quinolin-8-yl)azetidine-2-carboxamide (WX-04-500)
Prepared in similar fashion from *ent*-S2.
$^1$H NMR (400 MHz, $CD_3OD$): δ 8.90 – 8.83 (m, 1H), 8.29 – 8.24 (m, 1H), 8.20 (dd, *J* = 14.9, 7.7 Hz, 1H), 7.61 – 7.51 (m, 2H), 7.43 (q, *J* = 7.7 Hz, 1H), 7.35 (dd, *J* = 8.2, 3.9 Hz, 2H), 7.19 (d, *J* =

8.3 Hz, 2H), 5.62 (d, $J$ = 7.9 Hz, 0.55H), 5.40 (d, $J$ = 9.7 Hz, 0.45H), 4.69 – 4.57 (m, 1H), 4.49 – 4.34 (m, 2H), 3.34 (s, 1H), 2.12 (s, 1.35H), 1.83 (s, 1.65H); 1 exchangeable proton not observed; mixture of rotamers.
HRMS m/z calc. for $C_{25}H_{20}N_3O_2$ $[M+H]^+$ 394.1556 found 394.1559.

*Synthesis of WX-04-749 and WX-04-750*

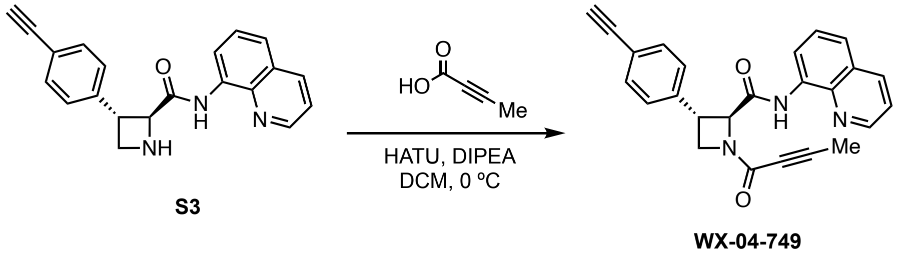


(2*S*,3*S*)-1-(but-2-ynoyl)-3-(4-ethynylphenyl)-*N*-(quinolin-8-yl)azetidine-2-carboxamide (WX-04-749)
To a precooled (0 °C) solution of but-2-ynoic acid (36.0 mg, 428 µmol, 2 equiv), HATU (163 mg, 428 µmol, 2 equiv) and DIPEA (82.9 mg, 641 µmol, 3 equiv) in DCM (2 mL) was added S3 (70 mg, 214 µmol, 1 equiv). The mixture was stirred at 0 °C for 1 h. Upon reaction completion, the mixture was concentrated under reduced pressure. The resulting residue was purified by prep-TLC ($SiO_2$, petroleum ether/EtOAc = 2:1) to obtain WX-04-749 (68.0 mg, 81% yield) as a yellow solid.
$^1$H NMR (400 MHz, $CD_3OD$): δ 8.91 – 8.79 (m, 1H), 8.78 – 8.67 (m, 1H), 8.32 (ddd, $J$ = 13.0, 8.3, 1.6 Hz, 1H), 7.74 – 7.38 (m, 7H), 5.30 (d, $J$ = 5.6 Hz, 0.45H), 5.12 (d, $J$ = 6.3 Hz, 0.55H), 4.66 (t, $J$ = 8.9 Hz, 0.55H), 4.53 (t, $J$ = 9.4 Hz, 0.45H), 4.36 (dd, $J$ = 8.9, 6.6 Hz, 0.55H), 4.16 (td, $J$ = 9.6, 5.9 Hz, 1H), 4.06 (q, $J$ = 6.3 Hz, 0.45H), 3.53 – 3.50 (m, 1H), 2.11 (s, 1.65H), 1.74 (s, 1.35H); 1 exchangeable proton not observed; mixture of rotamers.
HRMS m/z calc. for $C_{25}H_{20}N_3O_2$ $[M+H]^+$ 394.1556 found 394.1557.

(2*R*,3*R*)-1-(but-2-ynoyl)-3-(4-ethynylphenyl)-*N*-(quinolin-8-yl)azetidine-2-carboxamide (WX-04-750)
Prepared in similar fashion from *ent*-S3.
$^1$H NMR (400 MHz, $CD_3OD$): δ 8.87 (d, $J$ = 4.3 Hz, 1H), 8.73 (t, $J$ = 8.3 Hz, 1H), 8.32 (dd, $J$ = 12.4, 8.3 Hz, 1H), 7.75 – 7.35 (m, 7H), 5.31 (d, $J$ = 5.6 Hz, 0.45H), 5.12 (d, $J$ = 6.3 Hz, 0.55H), 4.66 (t, $J$ = 8.9 Hz, 0.55H), 4.53 (t, $J$ = 9.4 Hz, 0.45H), 4.40 – 4.32 (m, 0.55H), 4.16 (td, $J$ = 9.6, 6.4 Hz, 1H), 4.06 (q, $J$ = 6.1 Hz, 0.45H), 3.54 – 3.49 (m, 1H), 2.11 (s, 1.65H), 1.74 (s, 1.35H); 1 exchangeable proton not observed; mixture of rotamers.
HRMS m/z calc. for $C_{25}H_{20}N_3O_2$ $[M+H]^+$ 394.1556 found 394.1558.

*Synthesis of WX-02-24 and WX-02-44*

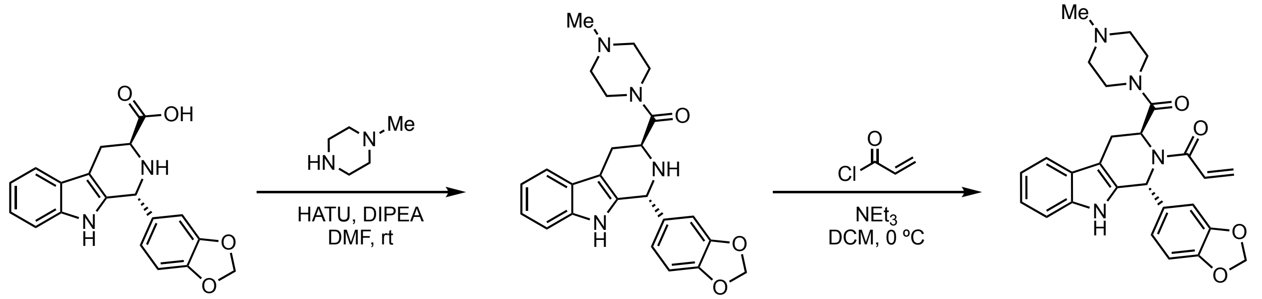


((1*R*,3*S*)-1-(benzo[*d*][1,3]dioxol-5-yl)-2,3,4,9-tetrahydro-1*H*-pyrido[3,4-*b*]indol-3-yl)(4-methylpiperazin-1-yl)methanone (S5)

To a solution of S4 (350 mg, 1.04 mmol, 1 equiv) in DMF (2 mL) were added HATU (594 mg, 1.56 mmol, 1.5 equiv), *N*-methyl piperazine (2.08 g, 20.8 mmol, 20 equiv) and DIPEA (269 mg, 2.08 mmol, 2 equiv). The mixture was stirred at rt for 1 h. Upon reaction completion, the mixture was diluted with water (20 mL) and extracted with EtOAc (20 mL × 3). The combined organic layers were dried over anhydrous sodium sulfate, filtered and concentrated under reduced. The resulting residue was purified by prep-HPLC (mobile phase: [A: water (10 mM $NH_4HCO_3$), B: acetonitrile]; gradient elution: 22%-52% B over 11 min) to obtain S5 (290 mg, 66% yield) as a white solid.

$^1$H NMR (400 MHz, $CDCl_3$) δ 7.86 (s, 1H), 7.55 (d, *J* = 7.7 Hz, 1H), 7.32 – 7.26 (m, 1H), 7.16 (dtd, *J* = 21.5, 7.1, 1.2 Hz, 2H), 6.76 – 6.69 (m, 2H), 6.63 (dd, *J* = 7.9, 1.8 Hz, 1H), 5.93 (s, 2H), 5.19 (s, 1H), 3.86 (dd, *J* = 10.2, 4.6 Hz, 1H), 3.78 – 3.65 (m, 1H), 3.60 – 3.47 (m, 1H), 3.32 – 3.23 (m, 1H), 3.23 – 3.12 (m, 1H), 3.04 (ddd, *J* = 15.8, 10.3, 1.4 Hz, 1H), 2.89 (dd, *J* = 15.7, 4.6 Hz, 1H), 2.49 – 2.21 (m, 8H); 2 exchangeable protons not observed.

LC-MS m/z calc. for $C_{24}H_{27}N_4O_3$ [M+H]$^+$ 419.2 found 419.2.

1-((1*R*,3*S*)-1-(benzo[*d*][1,3]dioxol-5-yl)-3-(4-methylpiperazine-1-carbonyl)-1,3,4,9-tetrahydro-2*H*-pyrido[3,4-*b*]indol-2-yl)prop-2-en-1-one (WX-02-24)

To a precooled (0 °C) solution of S5 (70.0 mg, 167 $\mu$mol, 1 equiv) in dichloromethane (2 mL) were added triethylamine (33.9 mg, 335 $\mu$mol, 2 equiv) and acryloyl chloride (15.1 mg, 167 $\mu$mol, 1 equiv). The mixture was stirred at 0 °C for 0.5 h. Upon reaction completion, the mixture was concentrated under reduced pressure. The resulting residue purified by prep-HPLC (mobile phase: [A: water (10 mM $NH_4HCO_3$), B: acetonitrile]; gradient elution: 26%-56% B over 10 min) to obtain WX-02-24 (39.5 mg, 49% yield) as a white solid.

$^1$H NMR (400 MHz, $CD_3OD$): δ 7.47 (d, *J* = 7.7 Hz, 1H), 7.27 (d, *J* = 8.0 Hz, 1H), 7.07 (ddt, *J* = 8.1, 7.0, 1.1 Hz, 1H), 7.01 (t, *J* = 7.4 Hz, 1H), 6.98 – 6.32 (m, 5H), 6.23 (d, *J* = 16.6 Hz, 1H), 5.92 (br. s, 2H), 5.74 (d, *J* = 10.5 Hz, 1H), 5.03 (br. s, 1H), 3.68 – 3.51 (m, 1H), 3.41 – 3.25 (m, 2H), 3.24 – 3.07 (m, 2H), 2.45 – 2.08 (m, 8H); 1 exchangeable proton not observed.

HRMS m/z calc. for $C_{27}H_{29}N_4O_4$ [M+H]$^+$ 473.2189 found 473.2200.

1-((1*S*,3*R*)-1-(benzo[*d*][1,3]dioxol-5-yl)-3-(4-methylpiperazine-1-carbonyl)-1,3,4,9-tetrahydro-2*H*-pyrido[3,4-*b*]indol-2-yl)prop-2-en-1-one (WX-02-44)

Prepared in similar fashion from *ent*-S4.

$^{1}H$ NMR (400 MHz, $CD_3OD$): δ 7.47 (d, *J* = 7.8 Hz, 1H), 7.27 (d, *J* = 8.0 Hz, 1H), 7.08 (t, *J* = 7.5 Hz, 1H), 7.01 (t, *J* = 7.5 Hz, 1H), 6.96 – 6.29 (m, 5H), 6.28 – 6.19 (m, 1H), 5.93 (br. s, 2H), 5.75 (d, *J* = 10.5 Hz, 1H), 5.04 (br. s, 1H), 3.70 – 3.52 (m, 1H), 3.42 – 3.30 (m, 2H), 3.25 – 3.07 (s, 2H), 2.44 – 2.11 (m, 8H); 1 exchangeable proton not observed.
HRMS m/z calc. for $C_{27}H_{29}N_4O_4$ $[M+H]^+$ 473.2189 found 473.2186.

#### Analytical data: NMR spectra

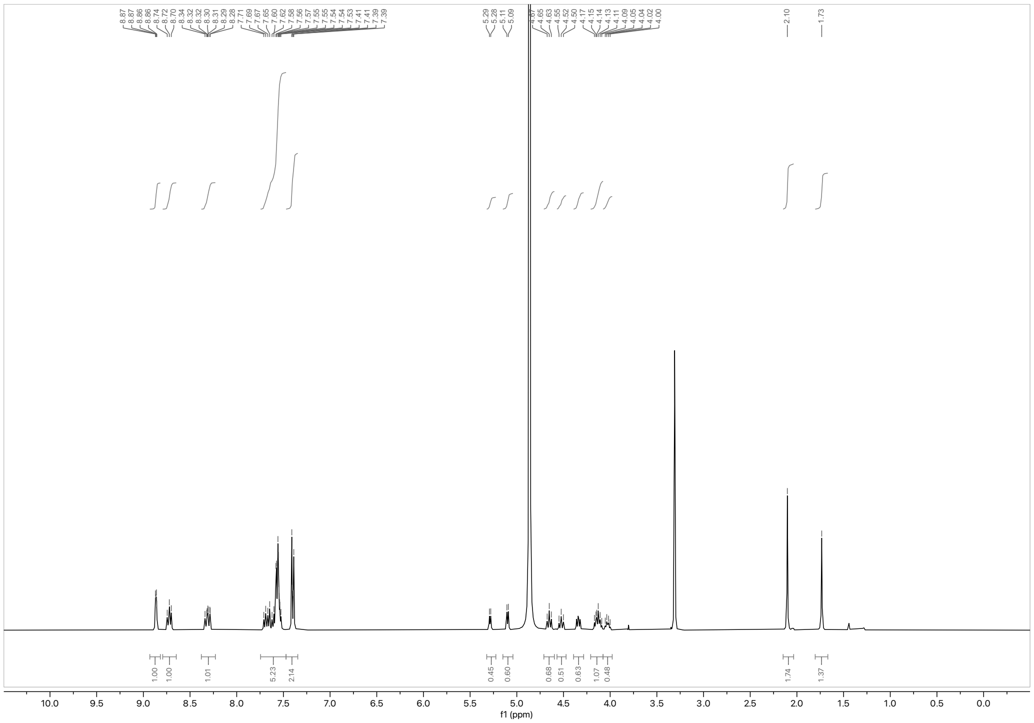

$^{1}H$ NMR spectrum of MY-46A (400 MHz, $CD_3OD$)

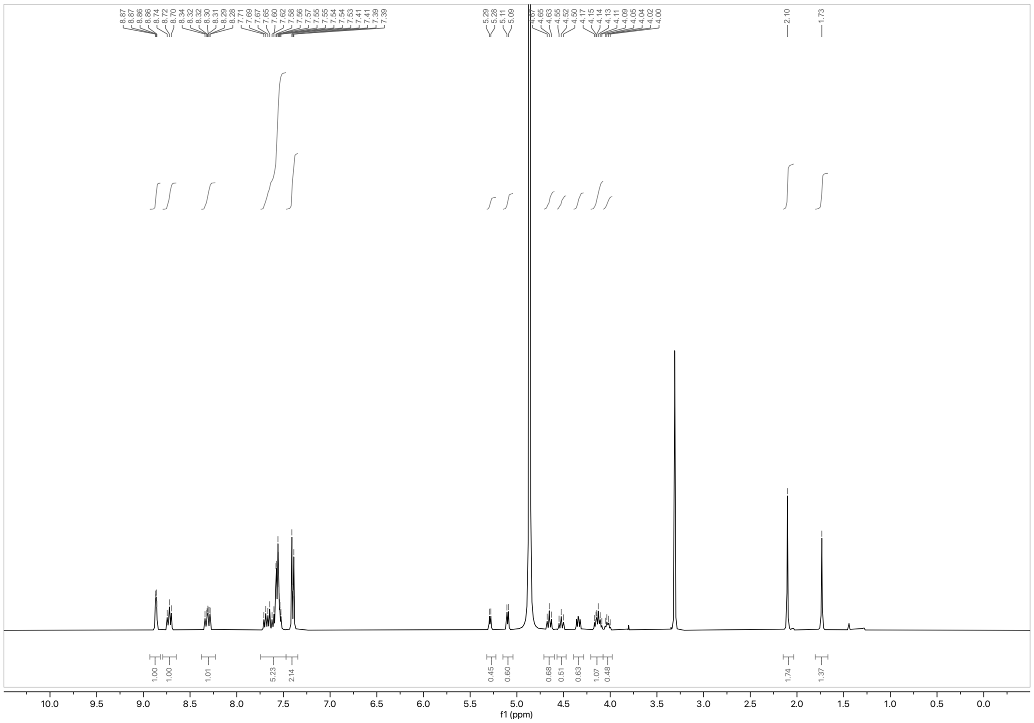

$^{1}H$ NMR spectrum of MY-46B (400 MHz, $CD_3OD$)

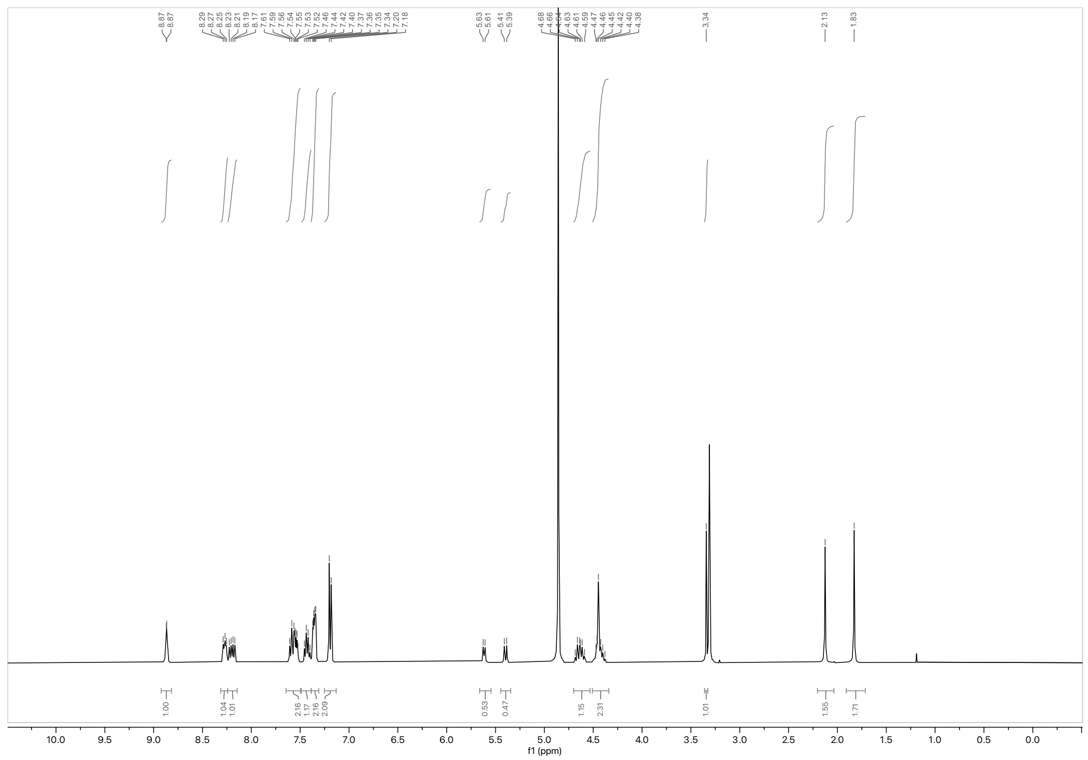

$^1H$ NMR spectrum of WX-04-499 (400 MHz, $CD_3OD$)

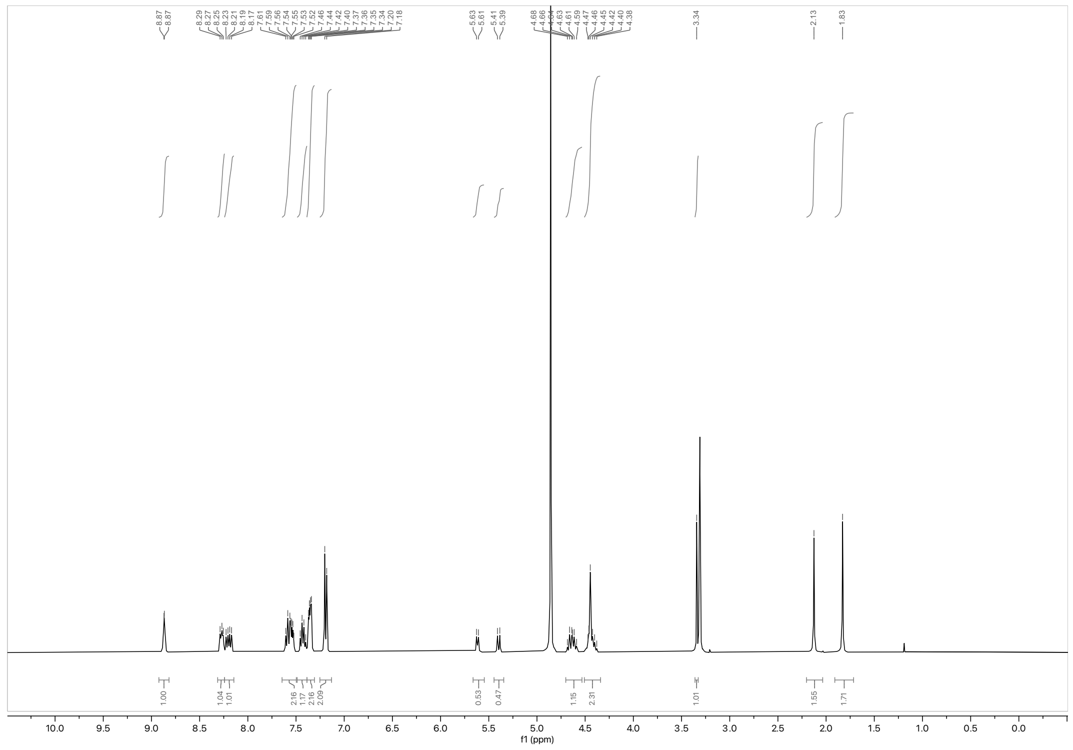

$^1H$ NMR spectrum of WX-04-500 (400 MHz, $CD_3OD$)

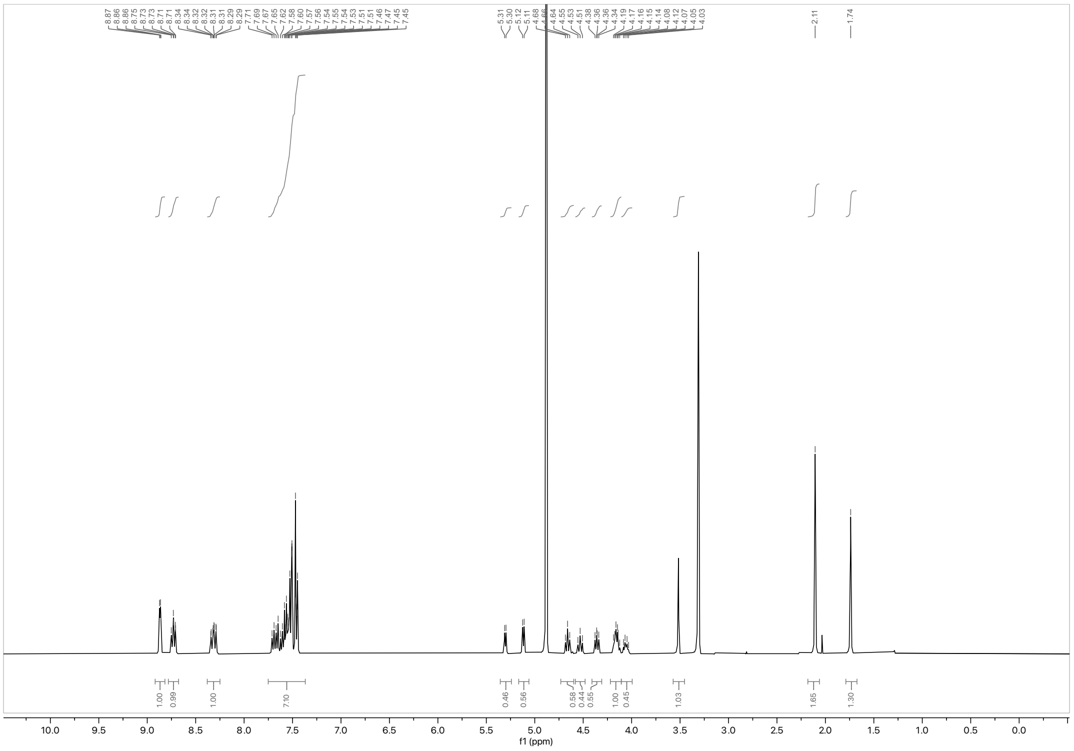

$^1H$ NMR spectrum of WX-04-749 (400 MHz, $CD_3OD$)

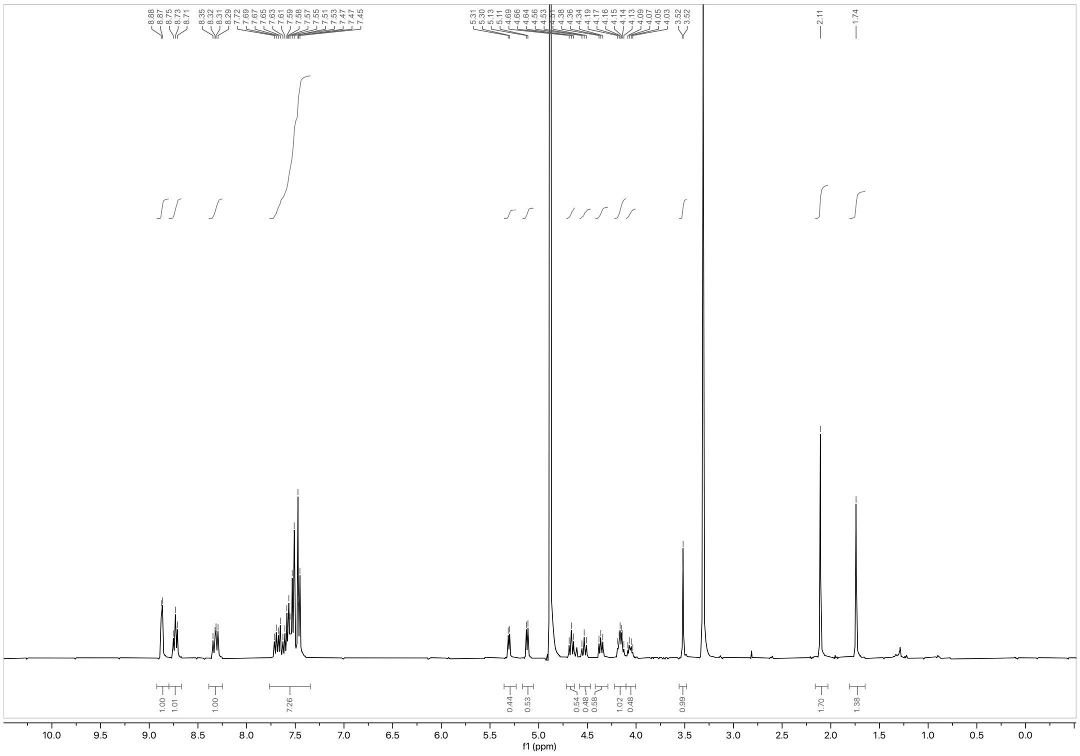

$^1H$ NMR spectrum of WX-04-750 (400 MHz, $CD_3OD$)

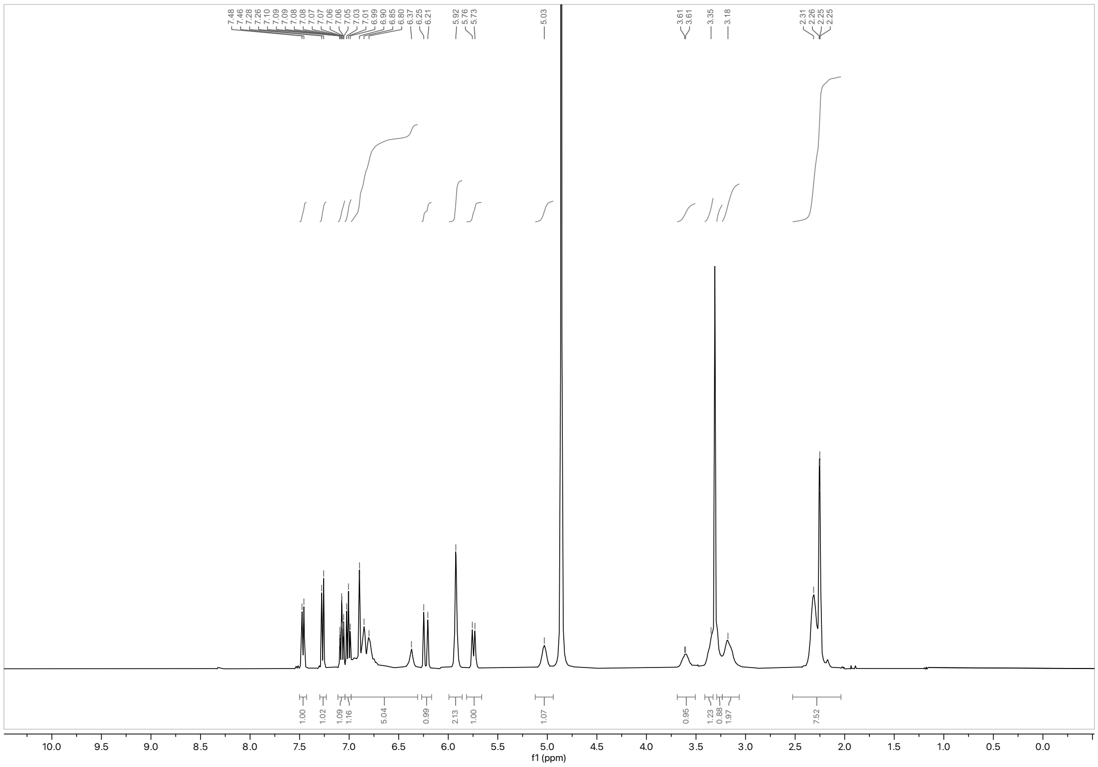

$^1H$ NMR spectrum of WX-02-24 (400 MHz, $CD_3OD$)

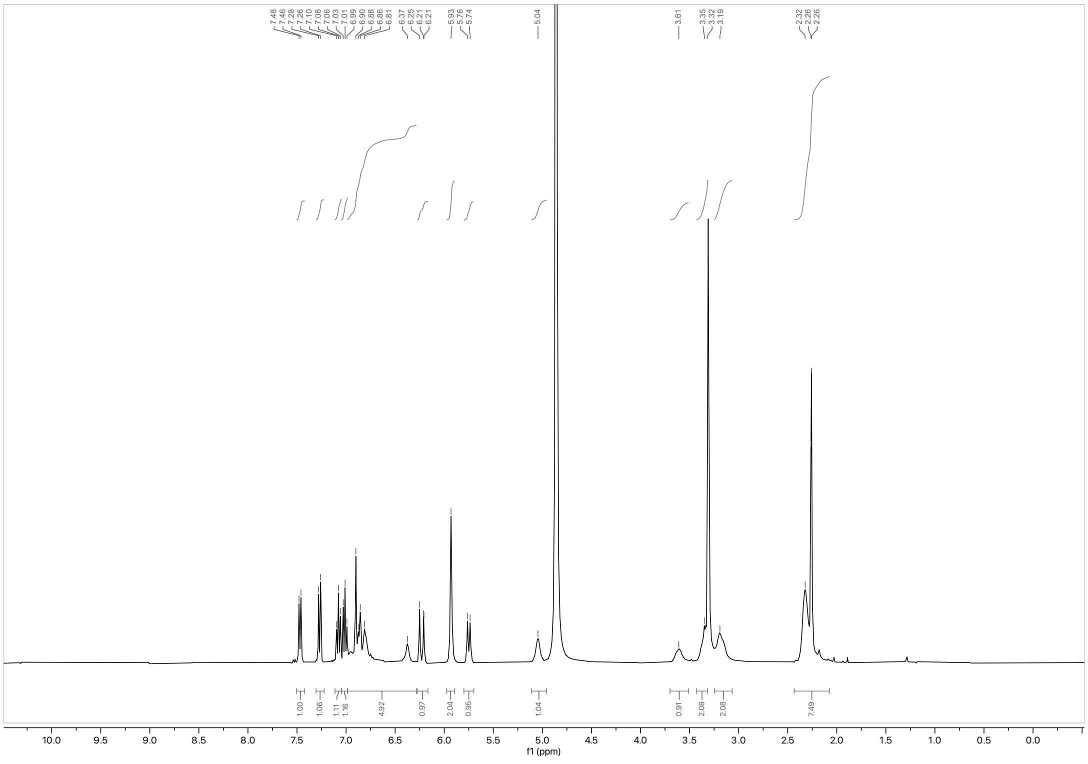

$^1H$ NMR spectrum of WX-02-44 (400 MHz, $CD_3OD$)

**Analytical data: SFC**

MY-46A

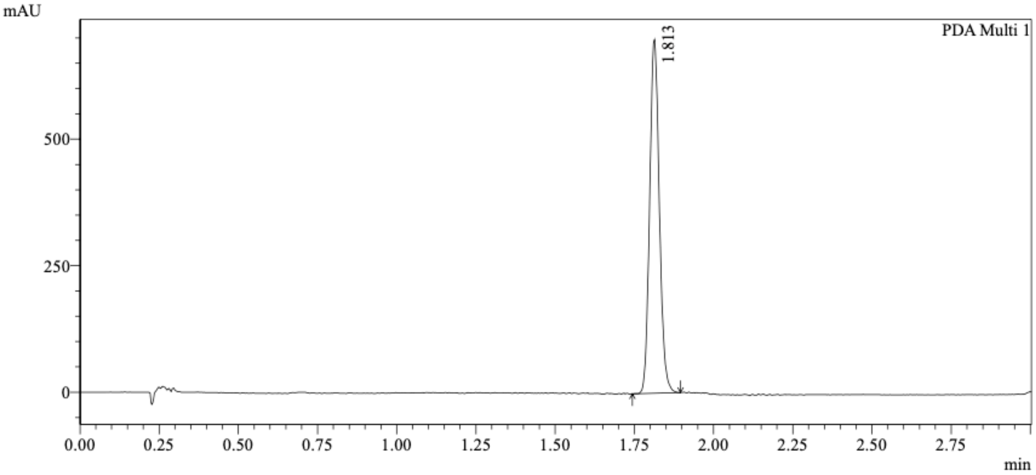


1 PDA Multi 1 / 220nm,4nm

Integration Results

PeakTable

PDA Ch1 220nm

| Peak# | Ret. Time | USP Width | Resolution | Height | Area | Area % |
|---|---|---|---|---|---|---|
| 1 | 1.813 | 0.059 | 0.000 | 689776 | 1482955 | 100.000 |
| Total | | | | 689776 | 1482955 | 100.000 |

MY-46B

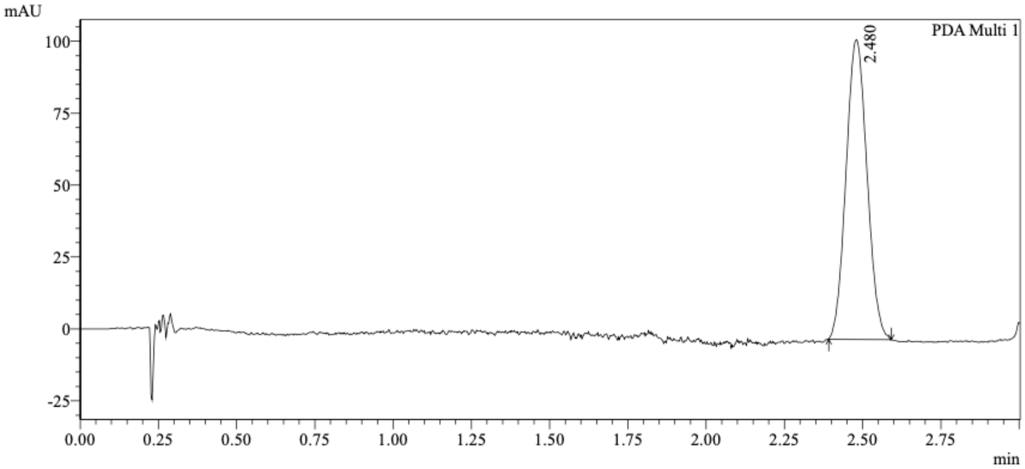


1 PDA Multi 1 / 220nm,4nm

Integration Results

PeakTable

PDA Ch1 220nm

| Peak# | Ret. Time | USP Width | Resolution | Height | Area | Area % |
|---|---|---|---|---|---|---|
| 1 | 2.480 | 0.127 | 0.000 | 103744 | 486225 | 100.000 |
| Total | | | | 103744 | 486225 | 100.000 |

Mixture of MY-46A & MY-45B

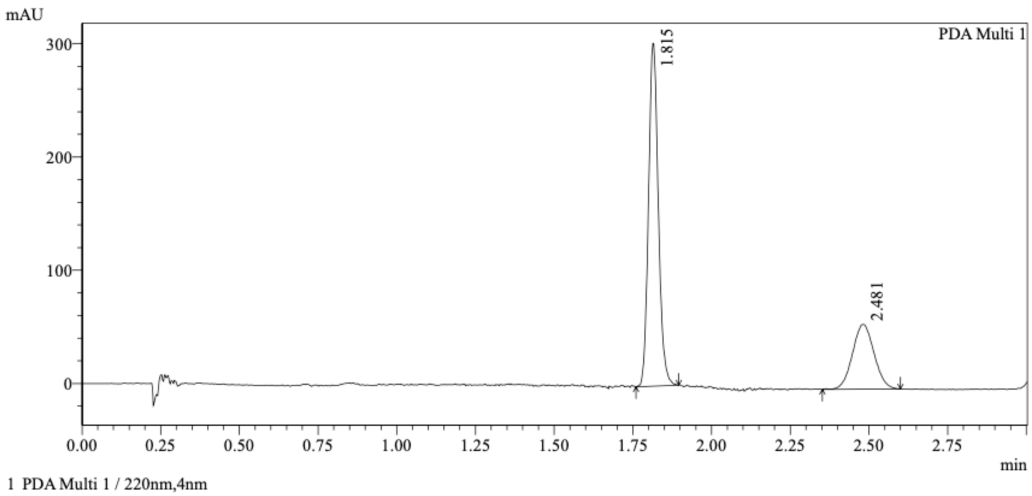


1 PDA Multi 1 / 220nm,4nm

Integration Results

PeakTable

PDA Ch1 220nm

| Peak# | Ret. Time | USP Width | Resolution | Height | Area | Area % |
|---|---|---|---|---|---|---|
| 1 | 1.815 | 0.060 | 0.000 | 299229 | 654165 | 70.593 |
| 2 | 2.481 | 0.128 | 7.088 | 56652 | 272512 | 29.407 |
| Total | | | | 355881 | 926677 | 100.000 |

Method details

Column: Chiralcel OJ-3 50×4.6mm I.D., 3 $\mu$m
Mobile phase: A: $CO_2$, B: MeOH (0.05% diethylamine)
Elution: 5%-40% B (gradient)

WX-04-499

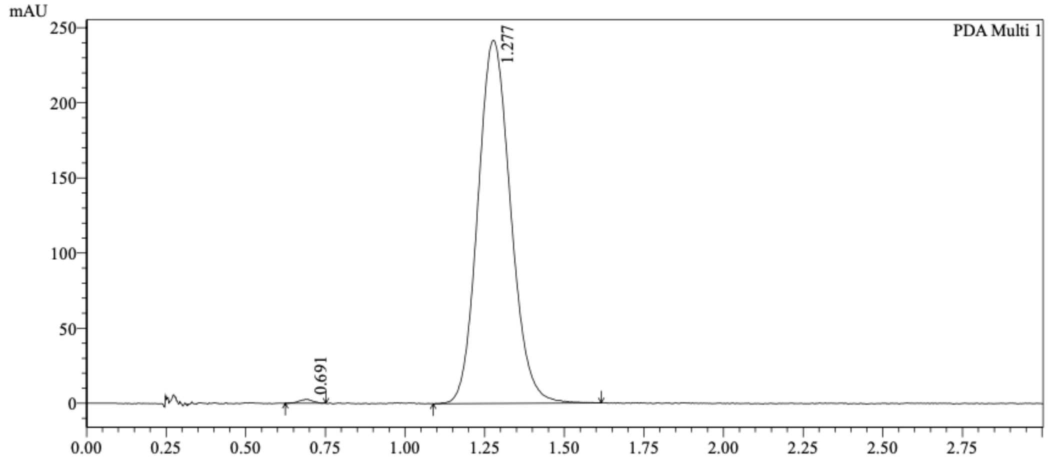


1 PDA Multi 1 / 220nm,4nm

Integration Results

PeakTable

PDA Ch1 220nm

| Peak# | Ret. Time | USP Width | Resolution | Height | Area | Area % |
|---|---|---|---|---|---|---|
| 1 | 0.691 | 0.086 | 0.000 | 2440 | 7500 | 0.432 |
| 2 | 1.277 | 0.188 | 4.281 | 241469 | 1728921 | 99.568 |
| Total | | | | 243909 | 1736422 | 100.000 |

WX-04-500

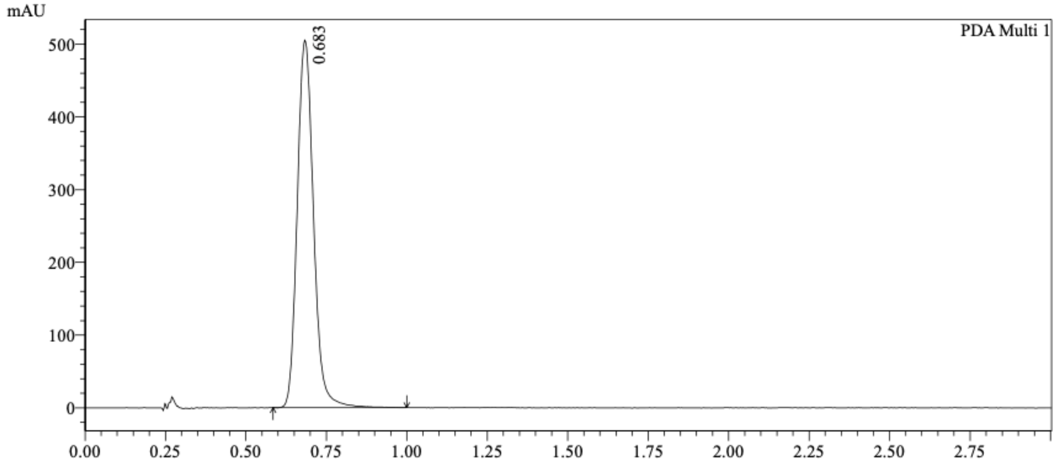


1 PDA Multi 1 / 220nm,4nm

Integration Results

PeakTable

PDA Ch1 220nm

| Peak# | Ret. Time | USP Width | Resolution | Height | Area | Area % |
|---|---|---|---|---|---|---|
| 1 | 0.683 | 0.093 | 0.000 | 501193 | 1776650 | 100.000 |
| Total | | | | 501193 | 1776650 | 100.000 |

Mixture of WX-04-499 & WX-04-500

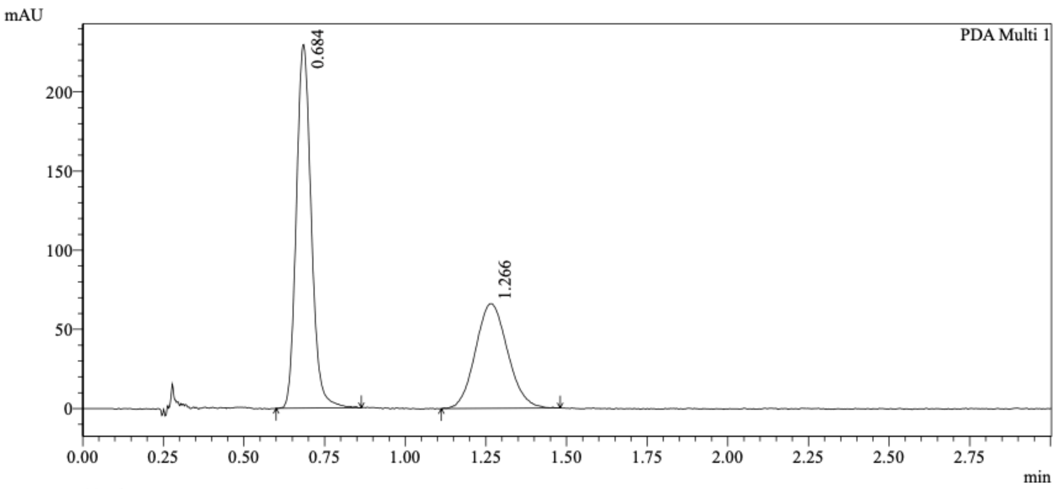


1 PDA Multi 1 / 220nm,4nm

Integration Results

PeakTable

PDA Ch1 220nm

| Peak# | Ret. Time | USP Width | Resolution | Height | Area | Area % |
|---|---|---|---|---|---|---|
| 1 | 0.684 | 0.086 | 0.000 | 228314 | 739498 | 61.702 |
| 2 | 1.266 | 0.184 | 4.316 | 65831 | 459005 | 38.298 |
| Total | | | | 294146 | 1198503 | 100.000 |

Method details

Column: Chiralpak AD-3 50×4.6mm I.D., 3 $\mu$m
Mobile phase: A: $CO_2$, B: MeOH (0.05% diethylamine)
Elution: 40% B

WX-04-749

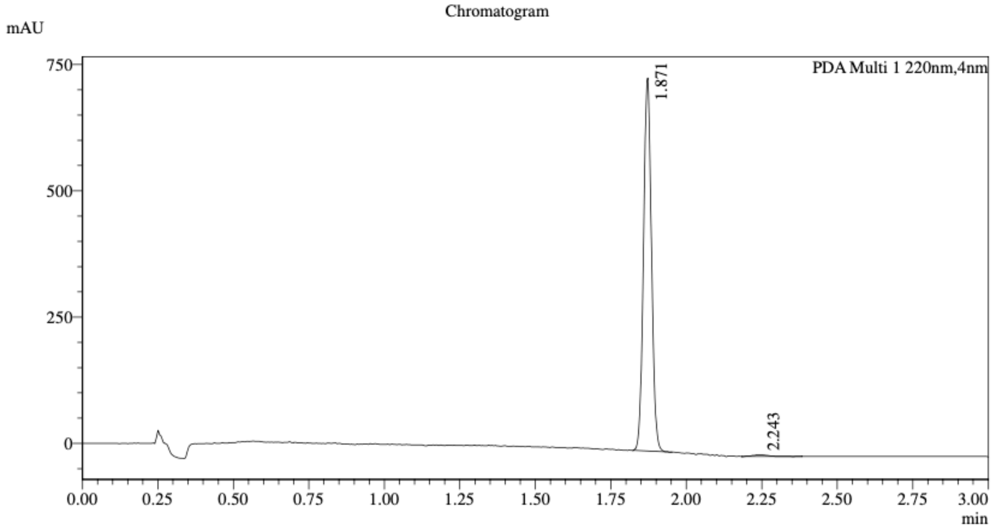


1 PDA Multi 1 / 220nm,4nm

Integration Result

Peak Table

PDA Ch1 220nm

| Peak# | Ret. Time | Height | Height% | Resolution(USP) | Area | Area% |
|---|---|---|---|---|---|---|
| 1 | 1.871 | 719265 | 99.532 | -- | 1353107 | 99.153 |
| 2 | 2.243 | 3380 | 0.468 | 5.031 | 11562 | 0.847 |

WX-04-750

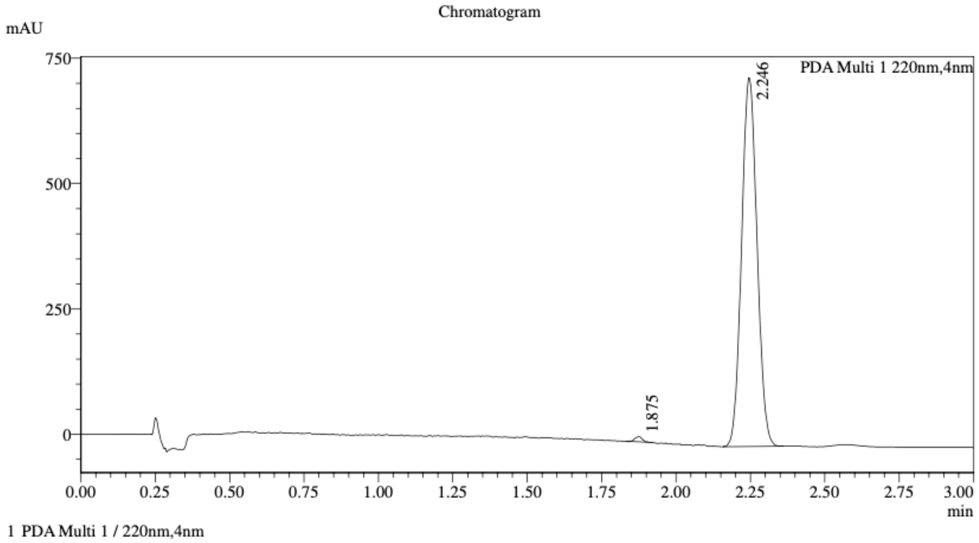


1 PDA Multi 1 / 220nm,4nm

Integration Result

Peak Table

PDA Ch1 220nm

| Peak# | Ret. Time | Height | Height% | Resolution(USP) | Area | Area% |
|---|---|---|---|---|---|---|
| 1 | 1.875 | 9990 | 1.345 | -- | 17458 | 0.653 |
| 2 | 2.246 | 732648 | 98.655 | 5.020 | 2657075 | 99.347 |

Mixture of WX-04-749 & WX-04-750

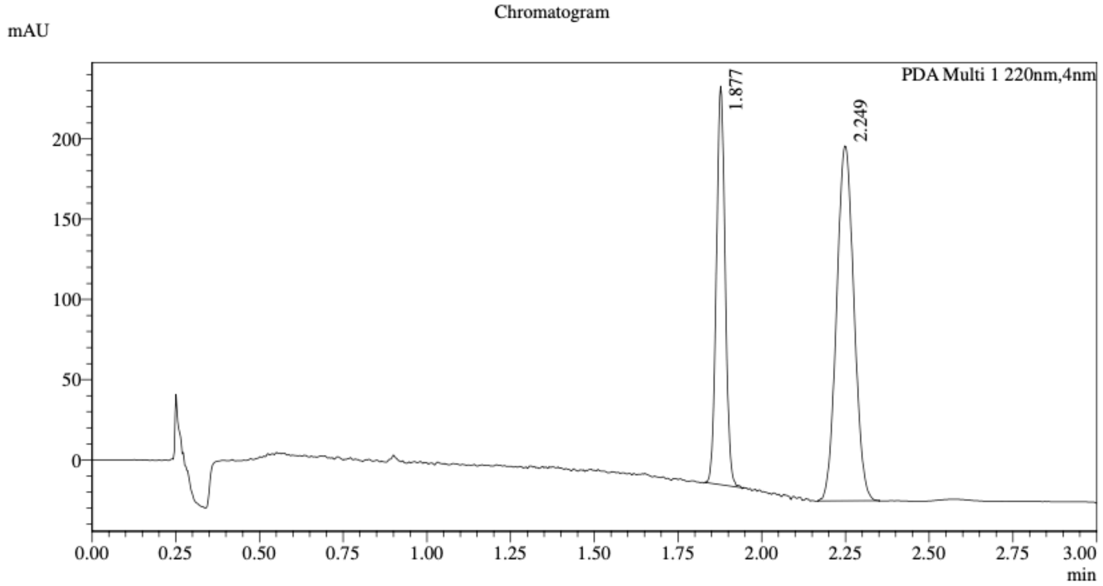


1 PDA Multi 1 / 220nm,4nm

Integration Result

Peak Table

PDA Ch1 220nm

| Peak# | Ret. Time | Height | Height% | Resolution(USP) | Area | Area% |
|---|---|---|---|---|---|---|
| 1 | 1.877 | 244241 | 52.862 | -- | 453703 | 36.272 |
| 2 | 2.249 | 217794 | 47.138 | 4.936 | 797143 | 63.728 |

Method details

Column: Chiralcel OJ-3 50×4.6mm I.D., 3 $\mu$m
Mobile phase: A: $CO_2$, B: MeOH (0.05% diethylamine)
Elution: 5%-40% B (gradient)

WX-02-24

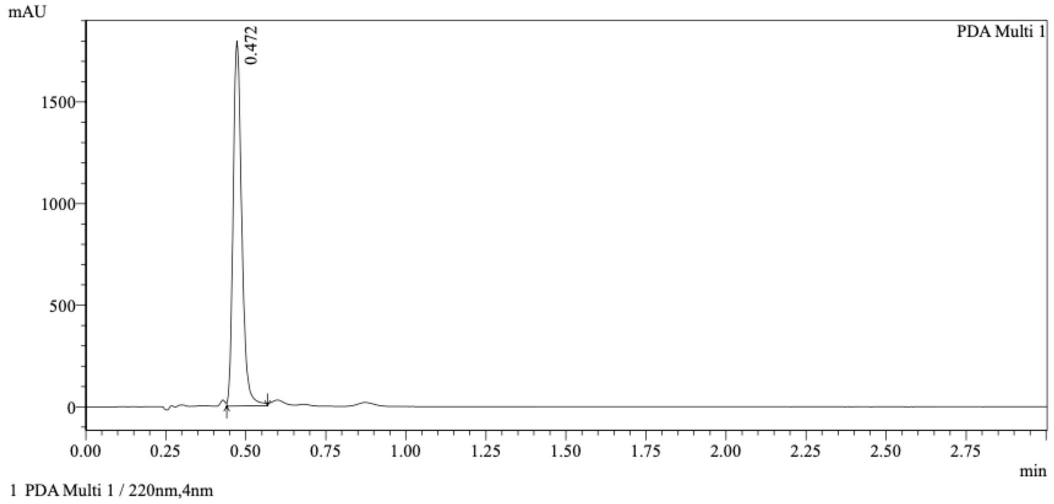


Integration Results

PeakTable

PDA Ch1 220nm

| Peak# | Ret. Time | USP Width | Resolution | Height | Area | Area % |
|---|---|---|---|---|---|---|
| 1 | 0.472 | 0.052 | 0.000 | 1711908 | 3360892 | 100.000 |
| Total | | | | 1711908 | 3360892 | 100.000 |

WX-02-44

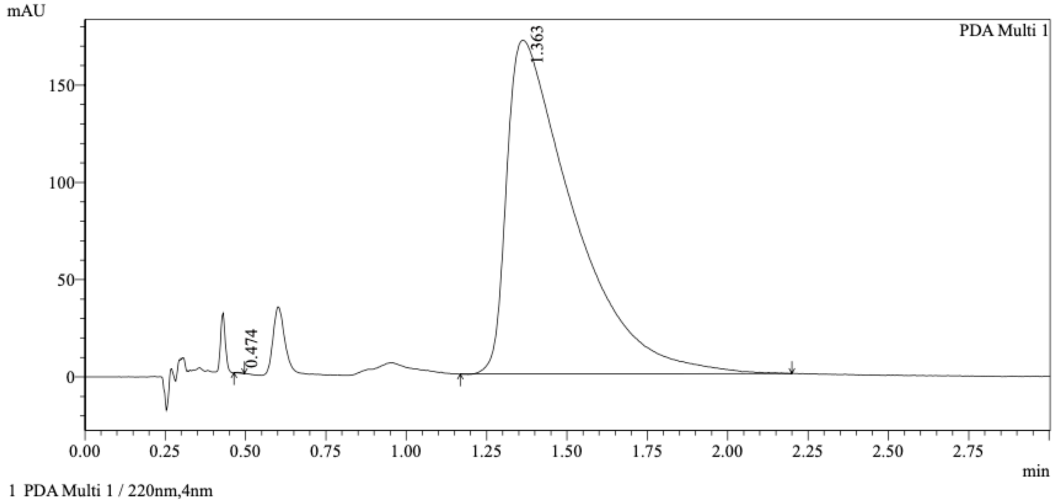


Integration Results

PeakTable

PDA Ch1 220nm

| Peak# | Ret. Time | USP Width | Resolution | Height | Area | Area % |
|---|---|---|---|---|---|---|
| 1 | 0.474 | 0.030 | 0.000 | 307 | 343 | 0.014 |
| 2 | 1.363 | 0.367 | 4.483 | 171367 | 2478824 | 99.986 |
| Total | | | | 171675 | 2479167 | 100.000 |

Mixture of WX-02-24 & WX-02-44

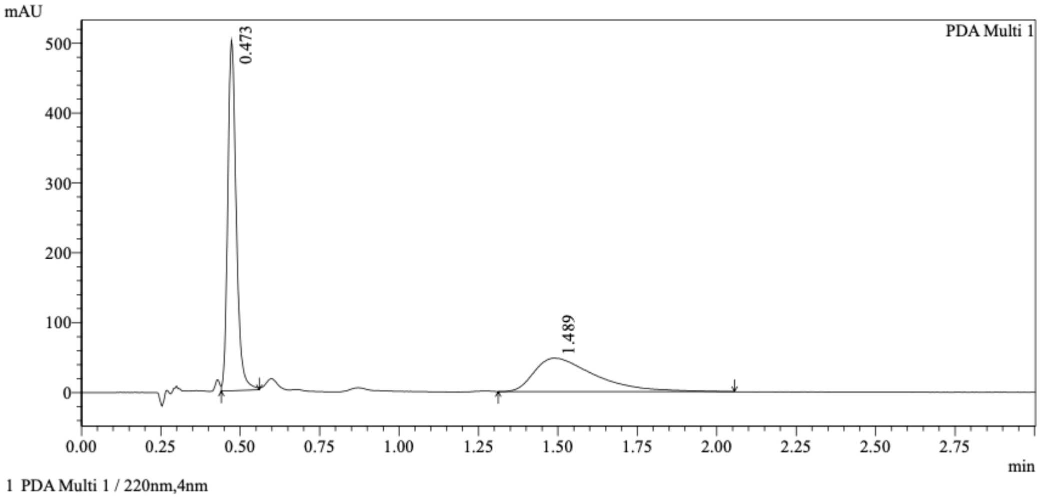


Integration Results

PeakTable

PDA Ch1 220nm

| Peak# | Ret. Time | USP Width | Resolution | Height | Area | Area % |
|---|---|---|---|---|---|---|
| 1 | 0.473 | 0.052 | 0.000 | 473639 | 931258 | 59.432 |
| 2 | 1.489 | 0.342 | 5.161 | 47705 | 635666 | 40.568 |
| Total | | | | 521345 | 1566925 | 100.000 |

Method details

Column: Chiralcel OD-3 50×4.6mm I.D., 3 $\mu$m
Mobile phase: A: $CO_2$, B: MeOH (0.05% diethylamine)
Elution: 40% B
